# Plasma amyloid β levels are driven by genetic variants near *APOE, BACE1, APP, PSEN2:* A genome-wide association study in over 12,000 non-demented participants

**DOI:** 10.1101/194266

**Authors:** Vincent Damotte, Sven J van der Lee, Vincent Chouraki, Benjamin Grenier-Boley, Jeannette Simino, Hieab Adams, Giuseppe Tosto, Charles White, Natalie Terzikhan, Carlos Cruchaga, Maria J. Knol, Shuo Li, Susanna Schraen, Megan L. Grove, Claudia Satizabal, Najaf Amin, Claudine Berr, Steven Younkin, Alzheimer’s Disease Neuroimaging Initiative, Rebecca F. Gottesman, Luc Buée, Alexa Beiser, David S. Knopman, Andre Uitterlinden, Charles DeCarli, Jan Bressler, Anita DeStefano, Jean-François Dartigues, Qiong Yang, Eric Boerwinkle, Christophe Tzourio, Myriam Fornage, M Arfan Ikram, Philippe Amouyel, Phil de Jager, Christiane Reitz, Thomas H Mosley, Jean-Charles Lambert, Sudha Seshadri, Cornelia van Duijn

**Author notes:** contributed equally to this work (Vincent Damotte, Sven J van der Lee, Vincent Chouraki). contributed equally to this work (Jean-Charles Lambert, Sudha Seshadri, Cornelia van Duijn). **Corresponding authors’ information**, Cornelia M van Duijn, Big Data Institute, Li Ka Shing Centre for Health Information and Discovery, Old Road Campus, Oxford, OX3 7LF, United Kingdom.

## Abstract

**INTRODUCTION:** There is increasing interest in plasma Aβ as an endophenotype and biomarker of Alzheimer’s disease (AD). Identifying the genetic determinants of plasma Aβ levels may elucidate important processes that determine plasma Aβ measures.

**METHODS:** We included 12,369 non-demented participants derived from eight population-based studies. Imputed genetic data and plasma Aβ1-40, Aβ1-42 levels and Aβ1-42/Aβ1-40 ratio were used to perform genome-wide association studies, gene-based and pathway analyses. Significant variants and genes were followed-up for the association with PET Aβ deposition and AD risk.

**RESULTS:** Single-variant analysis identified associations across *APOE* for Aβ1-42 and Aβ1-42/Aβ1-40 ratio, and *BACE1* for Aβ1-40. Gene-based analysis of Aβ1-40 additionally identified associations for *APP*, *PSEN2*, *CCK* and *ZNF397*. There was suggestive interaction between a *BACE1* variant and *APOE*ε4 on brain Aβ deposition.

**DISCUSSION:** Identification of variants near/in known major Aβ-processing genes strengthens the relevance of plasma-Aβ levels both as an endophenotype and a biomarker of AD.

## 1. Introduction

Aβ deposition is one of the hallmarks of Alzheimer’s disease (AD). Amyloid β (Aβ) peptides are the products of the catalytic processing of the Aβ precursor protein (APP) by the β-secretase, BACE1 and the γ-secretase complex [1]. Aβ peptides are able to self-assemble in soluble Aβ oligomers but also in insoluble fibrils that can aggregate as plaques in the brain parenchyma or in the wall of blood vessels where they constitute defining hallmarks of Alzheimer’s disease (AD) [2] and cerebral amyloid angiopathy (CAA), which is seen in many patients [3]. Aβ peptides are mainly produced in the brain where *APP* and *BACE1* are both highly expressed [1], but also in circulating blood platelets [4], in the pancreas [5] and the kidney [6].

There is strong evidence pointing toward a central role of Aβ peptides in the pathophysiology of AD [7]. Studies have shown that a large variety of rare mutations in genes involved in Aβ production, including *APP*, *PSEN1* and *PSEN2*, lead to autosomal dominant early-onset forms of AD and to lobar hemorrhage from cerebral amyloid angiopathy [8]. Moreover, apolipoprotein E (*APOE*) ε4, the major genetic risk factor for AD in the general population [9], has been implicated in Aβ aggregation, deposition and clearance, both in brain and in blood vessels [7, 10]. Although for long the Aβ –pathway did not emerge in our genome-wide association studies (GWAS) of AD [11], our most recent GWAS study highlighted Aβ –processing pathway and APP catabolic process pathway in late-onset Alzheimer’s disease (LOAD) [12]. We and others have also explored the genetics of Aβ through genome-wide association studies (GWAS) on quantitative measures of Aβ peptides, in the cerebrospinal fluid (CSF) or brain, through Pittsburgh Compound B (PiB) positron emission tomography (PET) scan or autopsy [13–17]. Combining the effect of AD genetic loci in a genetic risk score shows that the combined AD genes are statistically significantly related to CSF Aβ42 [17].

Although Aβ can be assessed in CSF and brain (PiB PET), these are of limited use for clinical and epidemiological studies in the population, either because of lower compliance (CSF) or higher costs (PiB PET). There is increasing interest in Aβ metabolism in blood. Aβ peptides produced in the brain can be degraded locally or transported into the CSF and the blood stream where they can be detected [18]. Although the brain-derived Aβ peptides in the circulation cannot be distinguished from Aβ derived from blood platelets, kidney or pancreas, a recent study using immunoprecipitation coupled with mass spectrometry to measure plasma Aβ1-40/Aβ1-42 and APP/Aβ1-42 ratios was able to accurately predict individual brain amyloid-β-positive or -negative status [19]. Also, studies assessing Aβ1-40 and Aβ1-42 using immunoassays show that these can predict Aβ status in the brain as assessed by PiB PET [20] and that changes in the blood and plasma occur simultaneously [21].

In our studies, we have also shown that plasma Aβ concentrations are prospectively associated with the future risk of developing AD [22–25]. Despite the fact that we have used less sensitive techniques to measure plasma Aβ levels, we found modest but significant correlation with amyloid burden in the CSF and in the brain [26, 27]. Identifying the genetic determinants of plasma Aβ levels may elucidate important processes that determine plasma Aβ measures. With this goal, we previously conducted a GWAS meta-analysis of plasma Aβ levels in 3,528 nondemented participants, but failed to find genome-wide significant associations [28], indicating a lack of power. As the more sensitive measures are not yet available in large samples with genome wide-genetic data we therefore aimed to increase the studied sample size of our previous work. The present study is a GWAS meta-analysis of plasma Aβ levels in over twelve thousands individuals aiming to elucidate processes that determine plasma beta amyloid.

## 2. Methods

### 2.1. Study populations

We included data from 12,369 European-descent participants from eight studies, the Framingham Heart Study (FHS; n=6,735), the Rotterdam study (RS, n=1,958), the Three City Study (3C; n=1,954), the Atherosclerosis Risk in Communities Study (ARIC; n=830), the Washington Heights-Inwood Community Aging Project (WHICAP; n=193), the Epidemiological Prevention study Zoetermeer (EPOZ; n=397), the Alzheimer’s Disease Neuroimaging Initiative (ADNI; n=173) and the Erasmus Rucphen Family Study (ERF; n=129). In each study, we excluded participants with prevalent dementia at the time of blood sampling used for plasma Aβ assessment (see Supplementary Materials and Methods 1 for a detailed description of each study).

### 2.2. Plasma Aβ assessment

Each study used different protocols for blood sampling, plasma extraction and storage and plasma Aβ assessment that have been detailed in previous publications [22, 23, 25, 29–31]. In the FHS, Rotterdam and 3C Study, plasma Aβ levels were measured at different times because of cost considerations. Various assays were used to quantify plasma Aβ1-40 and Aβ1-42 levels (see Supplementary Materials and Methods 2 for a detailed description of the protocols used in each study and Supplementary Table 1 for baseline characteristics of the study populations).

### 2.3. Genotyping

Each study used different genotyping platforms as previously published [11]. After applying preimputation variant and sample filters, genotypes were imputed using the 1000 Genomes phase 1 version 3 (all ethnicities) imputation panel and various imputation pipelines (see Supplementary Methods 3). *APOE* genotyping was performed as part of protocols specific to each study (see Supplementary Methods 4).

### 2.4. Statistical analyses

#### 2.4.1. Plasma Aβ levels

Plasma Aβ levels were expressed as pg/mL. In each study and for each Aβ dosage, we excluded values that were over or below 4 standard deviations around the mean. To study the variations of plasma Aβ levels in a consistent way across studies, we performed a ranked-based inverse normal transformation of plasma Aβ levels in each study. If they were significantly associated with plasma Aβ levels, this transformation was performed after adjusting for batch effect and other technical artifacts.

#### 2.4.2. Genome□wide association studies

Each study performed genome□wide association studies of plasma Aβ1-40 and Aβ1-42 levels and Aβ1-42/Aβ1-40 ratio using 1000 Genomes imputed data. According to the imputation pipelines used, genetic information was available either as allele dosages or genotype probabilities. In each study, we excluded results from variants that had low imputation quality (r2 or info score < 0.3), variants with low frequency (minor allele frequency < 0.005 or minor allele count < 7) and variants that were available in small number of participant (n < 30). Association of genetic variations with plasma Aβ levels were assessed in linear regression models adjusted for sex and age at blood collection. If significantly associated with plasma Aβ levels, principal components were added in the models to account for population structure.

#### 2.4.3. Genome□wide meta-analysis

Before meta-analysis, we applied a series of filters and quality check that were previously published (see Supplementary Figures 1 and 2) [32]. We performed an inverse variance weighted genome□wide meta-analysis, accounting for genomic inflation factors using the METAL software [33]. Finally, we retained variants that had been meta-analyzed at least in the 3 largest available populations (FHS, RS and 3C). Statistical significance was defined as a p-value below 5×10^-8^. Signals with p-values between 1×10^-5^ and 5×10^-8^ were considered suggestive. Additional graphs and analyses were done using R v3.6.1. To confirm the *APOE* signal we obtained in our genome□wide meta-analysis, we reran our analysis using genotyped *APOE* ε4 and *APOE* ε2 status, adjusting for age and sex.

#### 2.4.4. Gene-based and pathway analyses

We tested aggregated effects of SNPs located within genes using Multi-marker Analysis of GenoMic Annotation (MAGMA) v1.07 tool [34]. For each dosage, a total of 18,089 genes were tested, resulting in a significance threshold of 2.76×10^-6^. Pathway analyses were also performed with MAGMA v1.07 [34]. The following gene sets were used: GO (biological process, cellular component and molecular function, KEGG, Biocarta and Reactome). Pathway p-values were corrected for multiple testing using the FDR method.

#### 2.4.5. Association analyses with Aβ brain deposition

We related allelic variation at the SNP of interest with a standard measure of amyloid burden in the brain on Positron Emission Tomography (PET) imaging [35] in 193 Framingham Heart Study (FHS) participants [36] (see Supplementary Materials and Methods 5 for a detailed description of the protocols used). As a pre-specified hypothesis, we examined this association separately for persons with at least one *APOE* ε4 allele and those without. We report the odds ratio of having a positive amyloid scan associated with having a single copy of the allele of interest, using additive genetic models adjusted for age and sex.

#### 2.4.6. Association with AD

For significant variants and genes, we checked for association with AD. Summary statistics from the most recent genetic meta-analyses of AD were used [12, 37].

## 3. Results

### 3.1. Genome-wide significant variants associated with plasma Aβ levels

After meta-analysis, we identified 21 variants reaching genome-wide significance across two loci (Supplementary Figures 3 to 8).

The first locus was located on chromosome 19, in the *APOE* gene, with significant associations with plasma Aβ1-42 levels and plasma Aβ1-42/Aβ1-40 ratio (Figures 1 and 2). For both-13 – associations, the most significant variant was rs429358 with p-values of 9.01×10^-13^ and 6.46×10^-20^ for Aβ1-42 levels and Aβ1-42/Aβ1-40 ratio, respectively (Table 1). The minor allele of this variant, which denotes *APOE* ε4, was associated with lower plasma Aβ1-42 levels (effect size= −0.167 standard deviations (SD); 95% confidence interval (CI)=[−0.212; −0.121]) and lower plasma Aβ1-42/Aβ1-40 ratio (effect size=-0.212 SD; 95% CI=[−0.257; −0.121]; Table 1 and Supplementary Figure 9). We confirmed these associations using the directly genotyped *APOE* ε4 status (Supplementary Figure 10).

**Figure 1.**
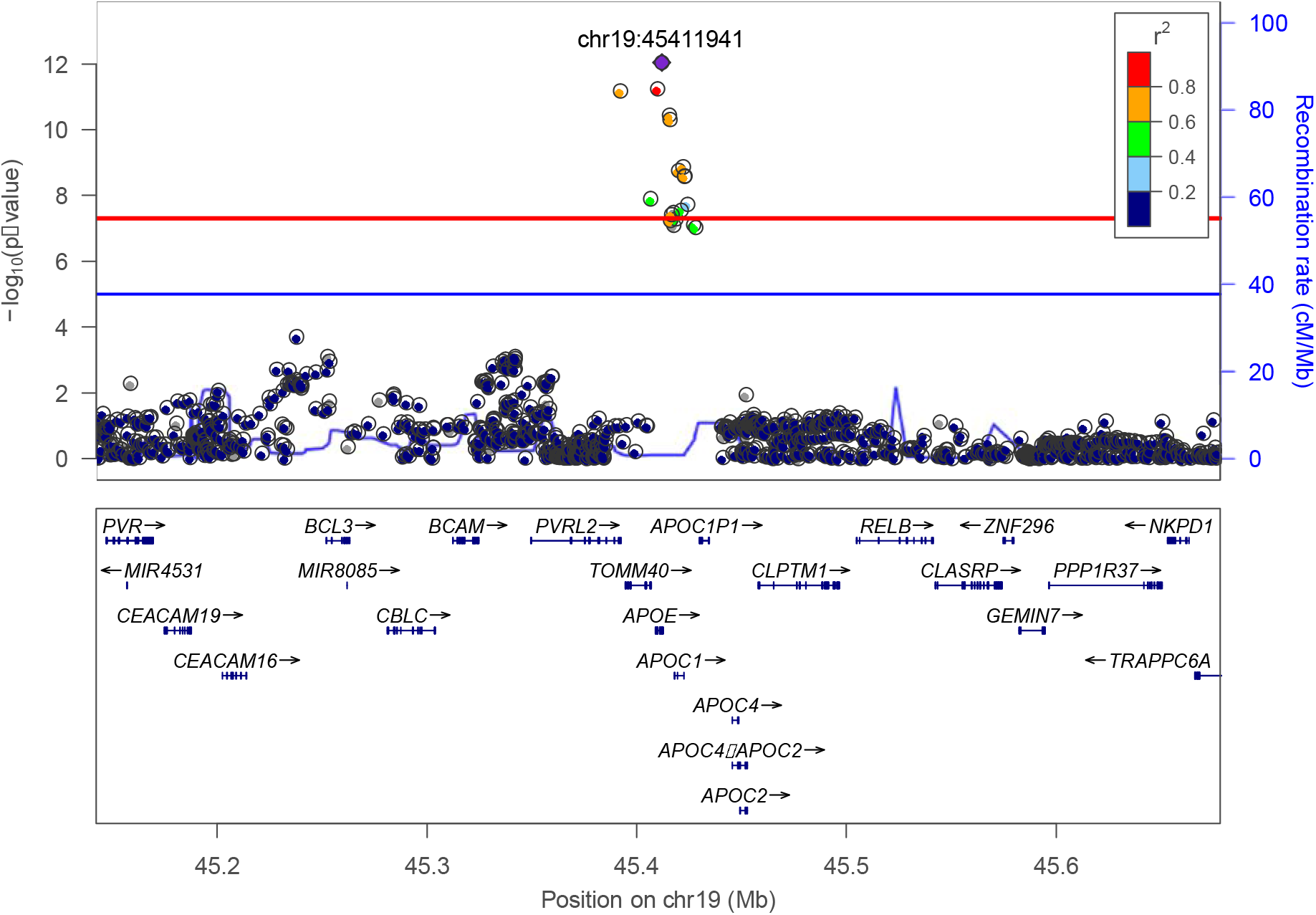
Association of frequent genetic variants with plasma Aβ1-42 in the *APOE* locus.

**Figure 2.**
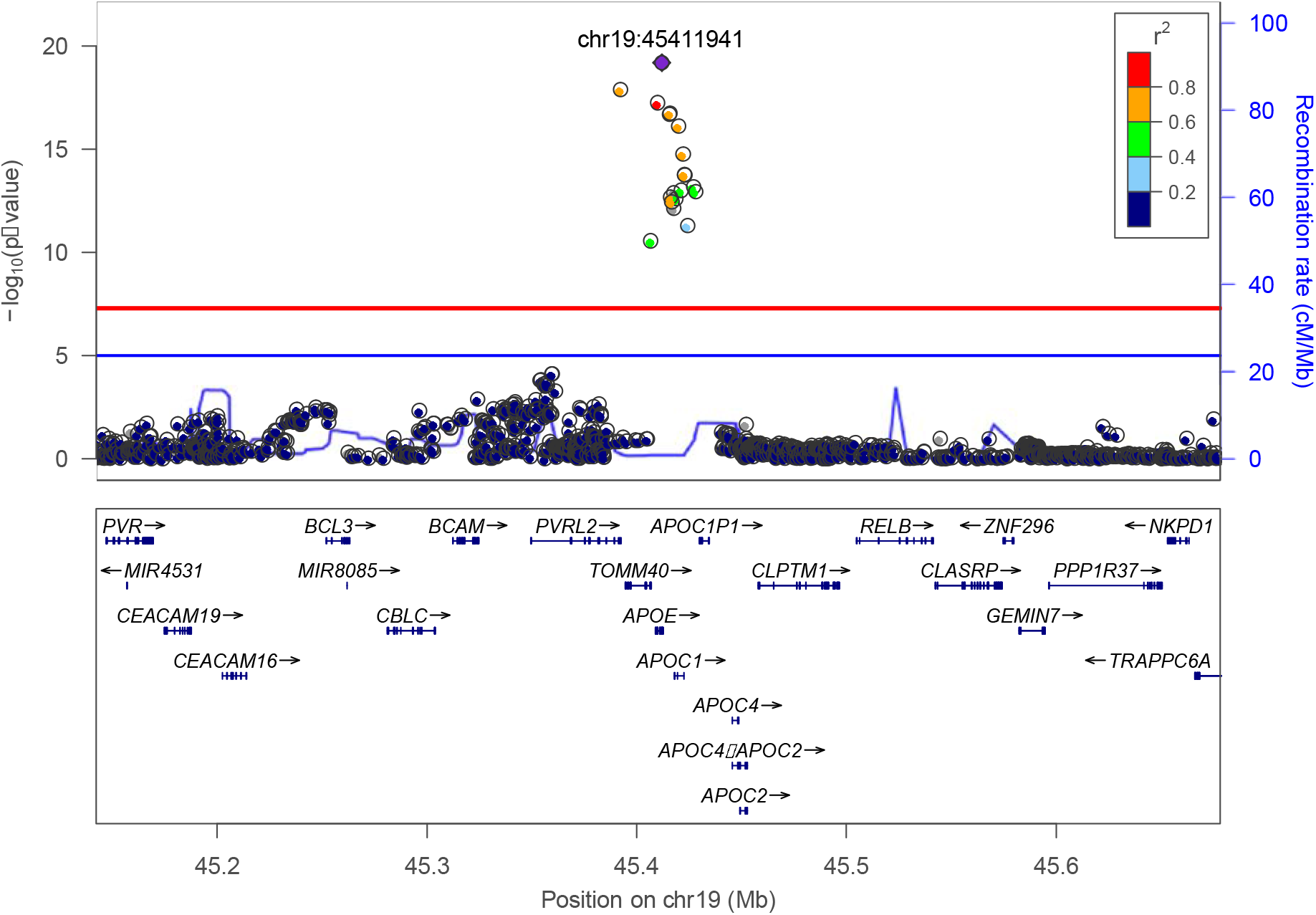
Association of frequent genetic variants with plasma Aβ1-42/Aβ1-40 ratio in the *APOE* locus.

**Table 1.** Association of top variants from genome-wide significant loci with plasma Aβ levels and amyloid-related traits.

|  | EAF | Effect | Standard Error | P-value | I<sup>2</sup> |
| --- | --- | --- | --- | --- | --- |
| **rs650585 (chr11:117110740, T/C, intron; RNF214/BACE1)** |  |  |  |  |  |
| *Plasma* |  |  |  |  |  |
| Aβ1-40 | 41.3% | -0.073 | 0.013 | 2.56x10<sup>-8</sup> | 3.1% |
| Aβ1-42 | 41.3% | -0.035 | 0.013 | 9.57x10<sup>-3</sup> | 27.8% |
| Aβ1-42/Aβ1-40 ratio | 41.4% | 0.033 | 0.013 | 1.39x10<sup>-2</sup> | 0.0% |
| *AD risk*<sup>a</sup> | 40.8% | 0.033 | 0.015 | 2.30x10<sup>-2</sup> | 8.3% |
| **rs429358 (chr19:45411941, C/T, missense; APOE)** |  |  |  |  |  |
| *Plasma* |  |  |  |  |  |
| Aβ1-40 | 13.4% | 0.023 | 0.023 | 3.11x10<sup>-1</sup> | 23.7% |
| Aβ1-42 | 13.4% | -0.167 | 0.023 | 9.01x10<sup>-13</sup> | 32.3% |
| Aβ1-42/Aβ1-40 ratio | 13.4% | -0.212 | 0.023 | 6.46x10<sup>-20</sup> | 52.6% |
| *AD risk*<sup>a</sup> | 21.6% | 1.20 | 0.019 | 0.00x10<sup>+00</sup> | 72.1% |

Abbreviations. EAF: effect allele frequency; AD: Alzheimer's Disease
Notes. For plasma measures, "Effect" represents the mean variation of the standardized variable (i.e. transformed so that mean=0 and standard deviation=1). In each block, the rsID of the top SNP is followed by its GRCh37 position, Effect/Non-effect alleles, functional category and closest genes. <sup>a</sup>: results obtained from [Kunkle\_2019]

The second genome□wide significant locus was an intronic variant in the *RNF214* gene. The function on *RNF214* is largely unknown. The gene is located on chromosome 11, near the *BACE1* gene. *BACE1* encodes the β-secretase and is involved in the initial, Aβ-producing step of APP processing (Figure 3). For the most significant variant, rs650585, the minor allele was associated with lower plasma Aβ1-40 levels (effect size=-0.073 SD; 95%CI=[−0.099; −0.047]; p-value=2.56×10^-8^; Table 1 and Supplementary Figure 9). This variant is in LD (R^2^=0.75, 1000 Genomes phase 3) with a *BACE1* synonymous variant, rs638405, which was also associated with plasma Aβ1-40 levels (effect size=-0.071 SD, p□value=1.21×10^-7^).

**Figure 3.**
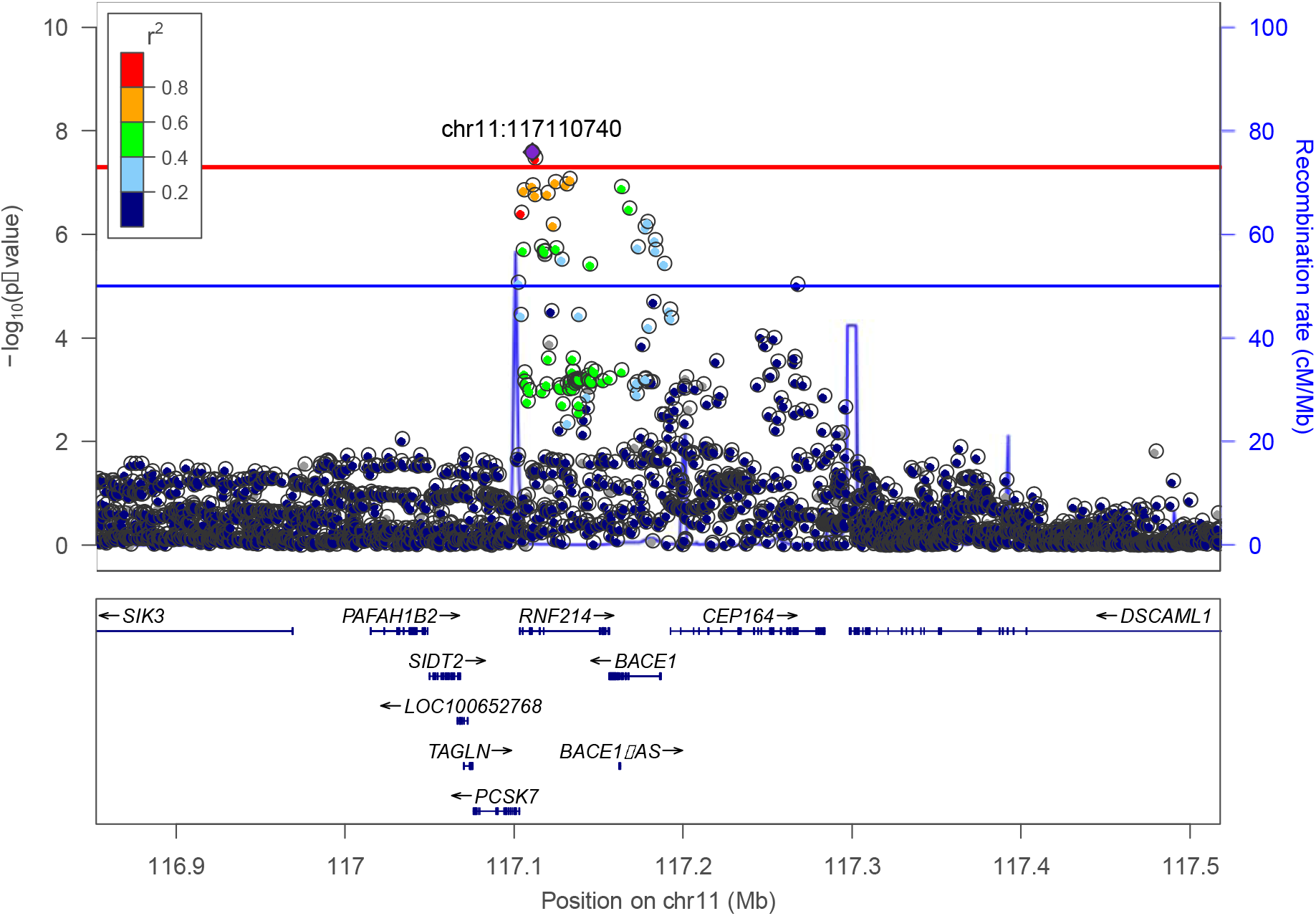
Association of frequent genetic variants with plasma Aβ1-40 in the BACE1 locus.

### 3.2. Gene and pathway-based analyses of plasma Aβ levels

Next, we performed gene-based tests (Table 2, Supplementary Figures 6-8). We again observed the *APOE, RNF214* and *BACE1* genes (p=3.87×10^-13^, p=2.33×10^-7^ and p=3.2×10^-9^, respectively), for which we had identified genome-wide significant single variant associations. Next to these genes, four genes showed gene-wide significant signals (p<2.76×10^-6^). We found that the *APP* and *PSEN2* genes were associated with plasma Aβ1-40 levels (p=1.67×10^-7^ and p=2.63×10^-6^, respectively). Interestingly, at the SNP level, there were two peaks reaching suggestive evidence for association with Aβ1-40 levels in *APP* gene (Supplementary Figure 11), probably explaining its strong association at the gene level. The two other genes were *CCK*, associated with plasma Aβ1-40 levels (p=2.63×10^-6^), and *ZNF397* associated with plasma Aβ1-42/1-40 ratio (p=2.27×10^-6^). The formal pathway analyses did not yield any significant results (Supplementary Tables 2-4).

**Table 2.**
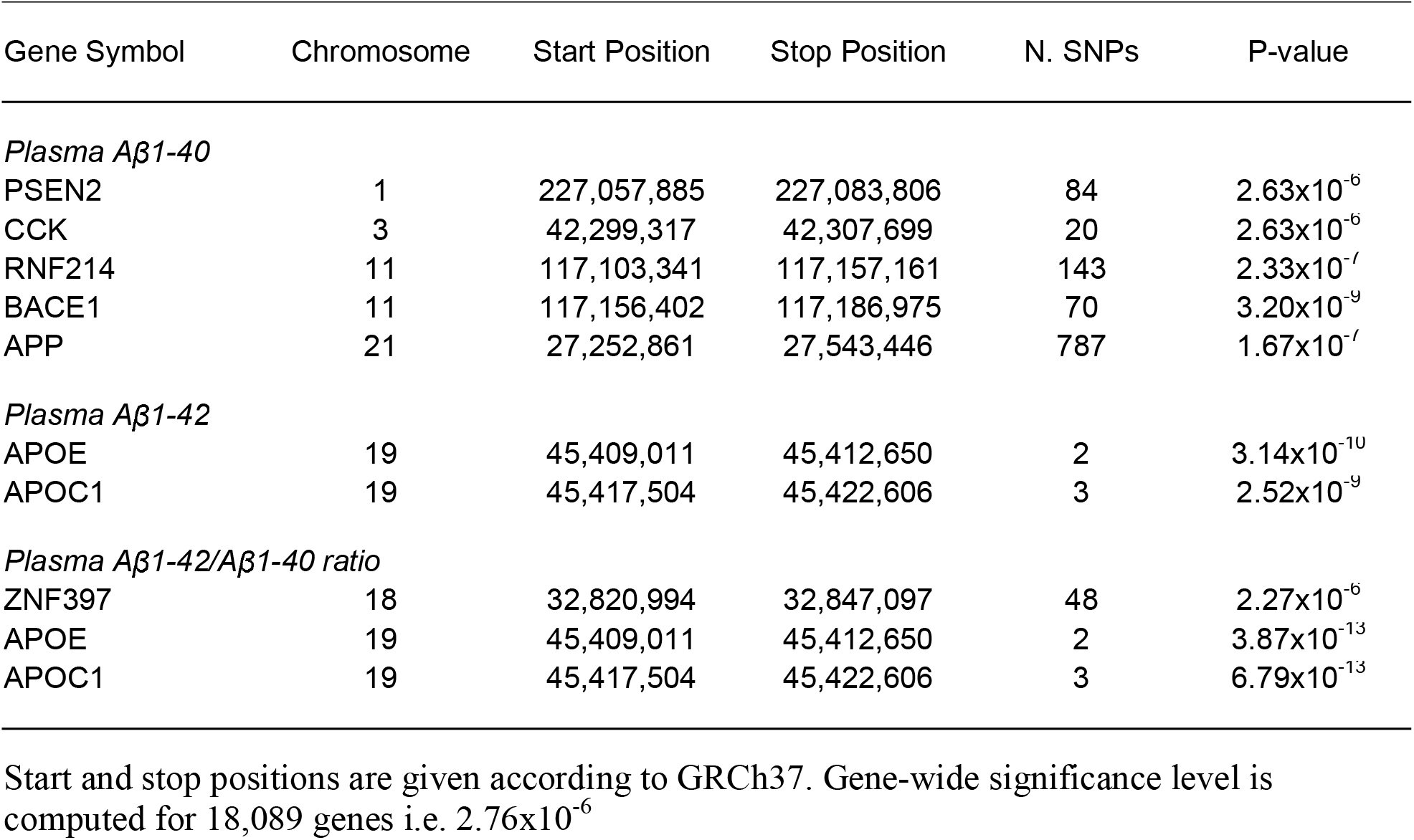
Associations of variants aggregated according to genes with plasma Aβ levels.

| Gene Symbol | Chromosome | Start Position | Stop Position | N. SNPs | P-value |
| --- | --- | --- | --- | --- | --- |
| *Plasma Aβ 1-40* |  |  |  |  |  |
| PSEN2 | 1 | 227,057,885 | 227,083,806 | 84 | 2.63x10<sup>-6</sup> |
| CCK | 3 | 42,299,317 | 42,307,699 | 20 | 2.63x10<sup>-6</sup> |
| RNF214 | 11 | 117,103,341 | 117,157,161 | 143 | 2.33x10<sup>-7</sup> |
| BACE1 | 11 | 117,156,402 | 117,186,975 | 70 | 3.20x10<sup>-9</sup> |
| APP | 21 | 27,252,861 | 27,543,446 | 787 | 1.67x10<sup>-7</sup> |
| *Plasma Aβ 1-42* |  |  |  |  |  |
| APOE | 19 | 45,409,011 | 45,412,650 | 2 | 3.14x10<sup>-10</sup> |
| APOC1 | 19 | 45,417,504 | 45,422,606 | 3 | 2.52x10<sup>-9</sup> |
| *Plasma Aβ 1-42/Aβ 1-40 ratio* |  |  |  |  |  |
| ZNF397 | 18 | 32,820,994 | 32,847,097 | 48 | 2.27x10<sup>-6</sup> |
| APOE | 19 | 45,409,011 | 45,412,650 | 2 | 3.87x10<sup>-13</sup> |
| APOC1 | 19 | 45,417,504 | 45,422,606 | 3 | 6.79x10<sup>-13</sup> |

Start and stop positions are given according to GRCh37. Gene-wide significance level is computed for 18,089 genes i.e. 2.76x10<sup>-6</sup>

### 3.3. Association of the BACE1 locus with PET Aβ deposition

We tested the association of the top hit rs650585 from the BACE1 locus (see above) with Aβ deposition in the brain from subsets of the FHS population. We found an association of rs650585 with an increase of deposition in FHS-Gen3 only among *APOE*ε4 positive individuals (p = 0.02) (Supplementary Table 5).

### 3.4. Variants associated with plasma amyloid associate with the risk of AD

The *APOE* □4 allele is known to be associated with a higher risk of AD [38]. We did not find significant evidence for association between genotyped *APOE* ε2 and circulating Aβ peptides levels, despite the protective effect of this variant on the risk of AD (Supplementary Figure 10).

A significant association of *APP* gene with AD (p=8.42×10^-7^) was reported [37]. Interestingly, one of the two peaks in *APP* reported as suggestively associated with Aβ1-40 levels (Supplementary Figure 11) was also associated with AD, whereas the second peak was not (Supplementary Figure 12) [37]. Nominal significant associations of *RNF214* (p=4.8×10^-5^) and *BACE1* (p=1.1×10^-3^) with AD were reported while *PSEN2* was close to nominal association (p=5.1×10^-2^) [37].

## 4. Discussion

Plasma Aβ1-40 and Aβ1-42 levels are increasingly of interest as biomarkers for AD. To uncover the important molecular processes underlying plasma Aβ we performed GWAS of plasma Aβ measured in 12,369 non□demented subjects. Despite that we did not use the recently developed specific immune-assays we uncovered that plasma Aβ is influenced by variants in and near *APOE*, *BACE1*, *PSEN2* and *APP*. These four genes code for known key proteins involved in Aβ processing. We also identified additional signals for the genes *CCK* and *ZNF397*. The variant near *BACE1* seemed also to be associated with Aβ in the brain as measured by PET imaging and were also associated with the risk of AD. In summary, plasma Aβ can be used to disentangle its molecular background.

The *BACE1* region encompasses several genes (*PCSK7*, *RNF214*, *BACE1*, *CEP164*) and a *BACE1* anti-sense long non-coding RNA (*BACE1*-*AS*). Although the top variant in the GWAS is located in an intron of *RNF214*, the gene-based analyses shows a significant association to *BACE1* that is more significant than the gene based test of RNF214. Since the β-secretase activity of BACE1 is necessary for Aβ peptide production, it is likely that *BACE1* or a local regulation of *BACE1* expression are responsible for this signal. We also found gene-wide significant associations with plasma Aβ1-40 levels in *APP* and *PSEN2*, two major actors of the Aβ metabolism. APP is obviously a central element of its own metabolism and PSEN2 is a key component of the γ-secretase which processes the APP C99 fragment into Aβ peptides [1]. The top variants at the *PSEN2* and *BACE1* loci were also nominally associated with Aβ1-42 levels in the same direction as Aβ1-40 levels, which is in agreement with knowledge that PSEN2 and BACE1 activities indifferently produce Aβ40 and Aβ42 peptides. Conversely, the *APOE*ε4 allele had the strongest association with Aβ1-42 levels but was not even nominally associated with Aβ1-40. This suggests that the APOE □4 isoform is not involved in the early process of Aβ peptide production but in more downstream events, such as Aβ aggregation or clearance. These results might also illustrate the greater ability to aggregate of Aβ1-42 peptides compared to Aβ1-40, and the influence of APOE isoforms in the regulation of this process [10]. Interestingly, associations of *APOE* ε2 with plasma Aβ levels were not significant and effect sizes were very small. Contrary to *APOE* ε4, the effect of *APOE* ε2 on amyloid markers has been much less studied and seems to be focused on specific brain regions, which could explain why we could not detect any association [39]. This could also suggest that other, Aβ□independent, mechanisms are involved in the lower risk of AD observed in *APOE* ε2 carriers [40].

The *CCK* gene is located in a region that was reported in a GWAS on neurofibrillary tangle [16]. Cholecystokinin (CCK) is a neuropeptide and gut hormone that regulates pancreatic enzyme secretion and gastrointestinal motility, and acts as a satiety signal. It is released simultaneously from intestinal cells and neurons in response to a meal. A sulfated form of cholecystokinin-8 may modulate neuronal activity in the brain [41]. The protein is located in axons, dendrites and the neuronal cell body and is involved in gastrin signaling and insulin secretion but also in neuron migration.

*ZNF397* gene encodes a protein with a N-terminal SCAN domain, and the longer isoform contains nine C2H2-type zinc finger repeats in the C-terminal domain. The protein localizes to centromeres during interphase and early prophase, and different isoforms can repress or activate transcription in transfection studies. Interestingly, the SNP rs509477, suggestively associated with CSF Aβ1-42 in a small association study [42], is located in an enhancer of *ZNF397* (Genecards: GH18J034976), acting in hippocampus middle, anterior caudate and cingulate gyrus brain regions [43]. However, this SNP was not associated with any of Aβ levels or ratio in our study.

Our analysis shows associations of plasma Aβ levels mainly with genes that are known for long to be involved in AD (*APOE*, *APP*, *PSEN2*), are nominally associated to AD or are expressed in brain regions. Although we cannot prove the origin, this suggests that Aβ peptides measured in the blood circulation probably originate from the brain rather than from the pancreas or the kidney. This perfectly fits with recent observations showing correlation of Aβ levels in blood with its levels in CSF as well as with its deposition in brain as assessed by PET imaging [19, 44].

For long, plasma Aβ is usually considered as a poor biomarker of AD in the literature. A previous meta-analysis reported that plasma Aβ levels were not useful to make a clinical diagnosis of AD [45]. Many of the cohorts participating in the present study have previously reported that low plasma Aβ42 and Aβ42/40 ratio levels were associated with development of AD after several years of follow-up [22–25]. The results of the present study are consistent with the hypothesis that Aβ in blood is predictive of AD pathophysiology and this view is strengthened by our present observation that *APOE* ε4, is both associated with low plasma Aβ42 and Aβ42/40 ratio and high AD risk. Some of those studies have also reported that this association remained significant after adjusting for *APOE* ε4 [25], suggesting that variations of plasma Aβ levels are not only an endophenotype of Aβ, but are also involved in AD pathophysiology. As such, plasma Aβ levels would not be only useful as a biomarker of an active amyloid metabolism in the brain but could also be considered as a biomarker for preventive interventions. In this light there are intriguing reports that hemodialysis or peritoneal dialysis are able to lower Aβ in the brain [46, 47]. Further, the association we observed between variants near *BACE1* and plasma Aβ40 is also of interest in the light of the ongoing trials testing BACE inhibitors, even though the lack of association of these variants with AD risk should be further investigated [48].

Our study has several strengths. First, it is, to date, the largest study of circulating amyloid peptides. This enabled us to identify factors of Aβ metabolism and we are optimistic about the relevance of the genetic signals that suggest blood levels of Aβ may have a clinical utility. Second, this study was conducted in non-demented participants and therefore is relevant for the study of early amyloid pathophysiological processes. Third, we carefully normalized the plasma Aβ data before running GWAS, thus taking into account some of the heterogeneity that has been described when using plasma Aβ levels.

Our study has also limitations. The state of current knowledge makes it difficult to extrapolate the role of these actors from the plasma compartments to the brain and further research in this area is needed. Second, the assays used in this study non-selectively measured Aβ concentrations and could not distinguish monomers from oligomers of Aβ, whether free or protein-bound. Therefore, our interpretation of the present results might differ from other studies in which assays used selectively measured monomers or oligomers of Aβ [49]. Future studies should carefully choose assays that allow measurements of each form of Aβ as this will facilitate interpretation with regard to the balance between Aβ production, aggregation and clearance.

In summary, our results indicate that genetic determinants of plasma Aβ40 and Aβ42 levels are close to genes known to be central actors in APP metabolism in AD. Increasing the statistical power of plasma Aβ analyses may potentially lead to the identification of currently unknown players in Aβ metabolism, novel hypotheses and hopefully, new preventive or therapeutic targets against Alzheimer’s disease.

## Supporting information

Supplementary Tables Part1

Supplementary Tables Part2

## Framingham Heart Study

This work was supported by the National Heart, Lung and Blood Institute’s Framingham Heart Study (contracts N01-HC-25195 and HHSN268201500001I). This study was also supported by grants from the National Institute on Aging: AG054076, U01-AG049505, and AG008122 (S. Seshadri). S.Seshadri, A.Beiser and Q.Yang were also supported by additional grants from the National Institute on Aging (R01AG049607, AG033193, AG033040) and the National Institute of Neurological Disorders and Stroke (R01-NS017950, R01-NS087541). The SHARe (SNP Health Association Resource) project was funded by the National Heart, Lung, and Blood Institute. The Linux Cluster for Genetic Analysis (LinGA-II) on which part of the FHS computations were performed was funded by the Robert Dawson Evans Endowment of the Department of Medicine at Boston University School of Medicine and Boston Medical Center.

## Rotterdam Study

The Rotterdam Study is funded by Erasmus Medical Center and Erasmus University, Rotterdam, Netherlands Organization for the Health Research and Development (ZonMw), the Research Institute for Diseases in the Elderly (RIDE), the Ministry of Education, Culture and Science, the Ministry for Health, Welfare and Sports, the European Commission (DG XII), and the Municipality of Rotterdam. The authors are grateful to the study participants, the staff from the Rotterdam Study and the participating general practitioners and pharmacists. The generation and management of the Illumina exome chip v1.0 array data for the Rotterdam Study (RS-I) was executed by the Human Genotyping Facility of the Genetic Laboratory of the Department of Internal Medicine (www.glimdna.org), Erasmus MC, Rotterdam, The Netherlands. The generation and management of GWAS genotype data for the Rotterdam Study (RS-I, RS-II, RS-III) was executed by the Human Genotyping Facility of the Genetic Laboratory of the Department of Internal Medicine, Erasmus MC, Rotterdam, The Netherlands. The GWAS datasets are supported by the Netherlands Organization of Scientific Research NWO Investments (nr. 175.010.2005.011, 911-03-012), the Genetic Laboratory of the Department of Internal Medicine, Erasmus MC, the Research Institute for Diseases in the Elderly (014-93-015; RIDE2), the Netherlands Genomics Initiative (NGI)/Netherlands Organization for Scientific Research (NWO) Netherlands Consortium for Healthy Aging (NCHA), project nr. 050-060-810. Carolina Medina-Gomez, PhD, Lennard Karsten, PhD, and Linda Broer, PhD, for QC and variant calling; Arp, Mila Jhamai, Marijn Verkerk, Lizbeth Herrera and Marjolein Peters, PhD, and Carolina Medina-Gomez, PhD, for their help in creating the GWAS database. The work for this manuscript was further supported by ADAPTED: Alzheimer’s Disease Apolipoprotein Pathology for Treatment Elucidation and Development (number 115975); the CoSTREAM project (www.costream.eu) and funding from the European Union’s Horizon 2020 research and innovation programme under grant agreement No 667375.

## Three City Study

This work was supported by INSERM, the National Foundation for Alzheimer’s disease and related disorders, the Institut Pasteur de Lille and the Centre National de Génotypage. This work has been developed and supported by the LABEX (laboratory of excellence program investment for the future) DISTALZ grant (Development of Innovative Strategies for a Transdisciplinary approach to Alzheimer’s disease) including funding from MEL (Metropoleeuropéenne de Lille), ERDF (European Regional Development Fund), Conseil Régional Nord Pas de Calais and the JPND-funded PERADES project. The Three-City Study was performed as part of collaboration between the Institut National de la Santé et de la Recherche Médicale (Inserm), the Victor Segalen Bordeaux II University and Sanofi-Synthélabo. The Fondation pour la Recherche Médicale funded the preparation and initiation of the study. The 3C Study was also funded by the Caisse Nationale Maladie des Travailleurs Salariés, Direction Générale de la Santé, MGEN, Institut de la Longévité, Agence Française de Sécurité Sanitaire des Produits de Santé, the Aquitaine and Bourgogne Regional Councils, Agence Nationale de la Recherche, ANR supported the COGINUT and COVADIS projects. Fondation de France and the joint French Ministry of Research/INSERM “Cohortes et collections de données biologiques” programme. Lille Génopôle received an unconditional grant from Eisai. The Three-city biological bank was developed and maintained by the laboratory for genomic analysis LAG-BRC - Institut Pasteur de Lille. The work for this manuscript was further supported by the CoSTREAM project (www.costream.eu) and funding from the European Union’s Horizon 2020 research and innovation programme under grant agreement No 667375.

## Atherosclerosis Risk in Communities Study

The Atherosclerosis Risk in Communities Study is carried out as a collaborative study supported by National Heart, Lung, and Blood Institute contracts (HHSN268201100005C, HHSN268201100006C, HHSN268201100007C, HHSN268201100008C, HHSN268201100009C, HHSN268201100010C, HHSN268201100011C, and HHSN268201100012C), R01HL087641, R01HL59367 and R01HL086694; National Human Genome Research Institute contract U01HG004402; and National Institutes of Health contract HHSN268200625226C. Neurocognitive data was collected by U01 HL096812, HL096814, HL096899, HL096902, and HL096917 from the NIH (NHLBI, NINDS, NIA and NIDCD), with previous brain MRI examinations funded by R01HL70825 from the NHLBI. Infrastructure was partly supported by Grant Number UL1RR025005, a component of the National Institutes of Health and NIH Roadmap for Medical Research. The authors thank the staff and participants of the ARIC study for their important contributions.

## Epidemiological Prevention study Zoetermeer

This research was made possible by financial support from the Netherlands Organization for Scientific Research and the Health Research Development Council.

## Washington Heights-Inwood Community Aging Project

Data collection and sharing for this project was supported by the Washington Heights-Inwood Columbia Aging Project (WHICAP, PO1AG007232, R01AG037212, RF1AG054023) funded by the National Institute on Aging (NIA) and by the National Center for Advancing Translational Sciences, National Institutes of Health, through Grant Number UL1TR001873. This work was also supported by National Institutes of Health grants AG042483, AG034189, and AG045334. This manuscript has been reviewed by WHICAP investigators for scientific content and consistency of data interpretation with previous WHICAP Study publications. We acknowledge the WHICAP study participants and the WHICAP research and support staff for their contributions to this study. The Columbia University Institutional Review Board reviewed and approved this project. All individuals provided written informed consent.

## Alzheimer’s Disease Neuroimaging Initiative

Data collection and sharing for this project was funded by the Alzheimer’s Disease Neuroimaging Initiative (ADNI) (National Institutes of Health Grant U01 AG024904) and DOD ADNI (Department of Defense award number W81XWH-12-2-0012). ADNI is funded by the National Institute on Aging, the National Institute of Biomedical Imaging and Bioengineering, and through generous contributions from the following: AbbVie, Alzheimer’s Association; Alzheimer’s Drug Discovery Foundation; Araclon Biotech; BioClinica, Inc.; Biogen; Bristol-Myers Squibb Company; CereSpir, Inc.; Cogstate; Eisai Inc.; Elan Pharmaceuticals, Inc.; Eli Lilly and Company; EuroImmun; F. Hoffmann-La Roche Ltd and its affiliated company Genentech, Inc.; Fujirebio; GE Healthcare; IXICO Ltd.; Janssen Alzheimer Immunotherapy Research & Development, LLC.; Johnson & Johnson Pharmaceutical Research & Development LLC.; Lumosity; Lundbeck; Merck & Co., Inc.; Meso Scale Diagnostics, LLC.; NeuroRx Research; Neurotrack Technologies; Novartis Pharmaceuticals Corporation; Pfizer Inc.; Piramal Imaging; Servier; Takeda Pharmaceutical Company; and Transition Therapeutics. The Canadian Institutes of Health Research is providing funds to support ADNI clinical sites in Canada. Private sector contributions are facilitated by the Foundation for the National Institutes of Health (www.fnih.org). The grantee organization is the Northern California Institute for Research and Education, and the study is coordinated by the Alzheimer’s Therapeutic Research Institute at the University of Southern California. ADNI data are disseminated by the Laboratory for Neuro Imaging at the University of Southern California.

## Eramus Rucphen Family Study

The Erasmus Rucphen Family (ERF) has received funding from the Centre for Medical Systems Biology (CMSB) and Netherlands Consortium for Systems Biology (NCSB), both within the framework of the Netherlands Genomics Initiative (NGI)/Netherlands Organization for Scientific Research (NWO). ERF study is also a part of EUROSPAN (European Special Populations Research Network) (FP6 STRP grant number 018947 (LSHG-CT-2006-01947)); European Network of Genomic and Genetic Epidemiology (ENGAGE) from the European Community’s Seventh Framework Programme (FP7/2007-2013)/grant agreement HEALTH-F4-2007-201413; “Quality of Life and Management of the Living Resources” of 5th Framework Programme (no. QLG2-CT-2002-01254); FP7 project EUROHEADPAIN (nr 602633), the Internationale Stichting Alzheimer Onderzoek (ISAO); the Hersenstichting Nederland (HSN); and the JNPD under the project PERADES (grant number 733051021, Defining Genetic, Polygenic and Environmental Risk for Alzheimer’s Disease using multiple powerful cohorts, focused Epigenetics and Stem cell metabolomics). This work in the ERF study was conducted under the grants: ADAPTED: Alzheimer’s Disease Apolipoprotein Pathology for Treatment Elucidation and Development (number 115975); the CoSTREAM project (www.costream.eu) and has received funding from the European Union’s Horizon 2020 research and innovation programme under grant agreement No 667375. We are grateful to all study participants and their relatives, general practitioners and neurologists for their contributions and to P. Veraart for her help in genealogy, J. Vergeer for the supervision of the laboratory work, and P. Snijders M.D. for his help in data collection.

The authors have no conflicts of interest to disclose.

## References

[1] Haass C, Kaether C, Thinakaran G, Sisodia S. Trafficking and proteolytic processing of APP. Cold Spring Harbor perspectives in medicine. 2012;2:a006270.

[2] Hyman BT, Phelps CH, Beach TG, Bigio EH, Cairns NJ, Carrillo MC, et al. National Institute on Aging-Alzheimer’s Association guidelines for the neuropathologic assessment of Alzheimer’s disease. Alzheimer’s & dementia: the journal of the Alzheimer’s Association. 2012;8:1–13.

[3] Viswanathan A, Greenberg SM. Cerebral amyloid angiopathy in the elderly. Annals of neurology. 2011;70:871–80.

[4] Chen M, Inestrosa NC, Ross GS, Fernandez HL. Platelets are the primary source of amyloid beta-peptide in human blood. Biochemical and biophysical research communications. 1995;213:96–103.

[5] Kulas JA, Puig KL, Combs CK. Amyloid precursor protein in pancreatic islets. The Journal of endocrinology. 2017;235:49–67.

[6] Selkoe DJ, Podlisny MB, Joachim CL, Vickers EA, Lee G, Fritz LC, et al. Beta-amyloid precursor protein of Alzheimer disease occurs as 110- to 135-kilodalton membrane-associated proteins in neural and nonneural tissues. Proceedings of the National Academy of Sciences of the United States of America. 1988;85:7341–5.

[7] Long JM, Holtzman DM. Alzheimer Disease: An Update on Pathobiology and Treatment Strategies. Cell. 2019;179:312–39.

[8] Cacace R, Sleegers K, Van Broeckhoven C. Molecular genetics of early-onset Alzheimer’s disease revisited. Alzheimer’s & dementia: the journal of the Alzheimer’s Association. 2016;12:733–48.

[9] Genin E, Hannequin D, Wallon D, Sleegers K, Hiltunen M, Combarros O, et al. APOE and Alzheimer disease: a major gene with semi-dominant inheritance. Molecular psychiatry. 2011;16:903–7.

[10] Kanekiyo T, Xu H, Bu G. ApoE and Abeta in Alzheimer’s disease: accidental encounters or partners? Neuron. 2014;81:740–54.

[11] Lambert JC, Ibrahim-Verbaas CA, Harold D, Naj AC, Sims R, Bellenguez C, et al. Metaanalysis of 74,046 individuals identifies 11 new susceptibility loci for Alzheimer’s disease. Nature genetics. 2013;45:1452–8.

[12] Kunkle BW, Grenier-Boley B, Sims R, Bis JC, Damotte V, Naj AC, et al. Genetic metaanalysis of diagnosed Alzheimer’s disease identifies new risk loci and implicates Abeta, tau, immunity and lipid processing. Nature genetics. 2019;51:414–30.

[13] Cruchaga C, Kauwe JS, Harari O, Jin SC, Cai Y, Karch CM, et al. GWAS of cerebrospinal fluid tau levels identifies risk variants for Alzheimer’s disease. Neuron. 2013;78:256–68.

[14] Ramanan VK, Risacher SL, Nho K, Kim S, Swaminathan S, Shen L, et al. APOE and BCHE as modulators of cerebral amyloid deposition: a florbetapir PET genome-wide association study. Molecular psychiatry. 2014;19:351–7.

[15] Shulman JM, Chen K, Keenan BT, Chibnik LB, Fleisher A, Thiyyagura P, et al. Genetic susceptibility for Alzheimer disease neuritic plaque pathology. JAMA neurology. 2013;70:1150–7.

[16] Beecham GW, Hamilton K, Naj AC, Martin ER, Huentelman M, Myers AJ, et al. Genomewide association meta-analysis of neuropathologic features of Alzheimer’s disease and related dementias. PLoS genetics. 2014;10:e1004606.

[17] Deming Y, Li Z, Kapoor M, Harari O, Del-Aguila JL, Black K, et al. Genome-wide association study identifies four novel loci associated with Alzheimer’s endophenotypes and disease modifiers. Acta neuropathologica. 2017;133:839–56.

[18] Sagare AP, Bell RD, Zlokovic BV. Neurovascular defects and faulty amyloid-beta vascular clearance in Alzheimer’s disease. Journal of Alzheimer’s disease: JAD. 2013;33 Suppl 1:S87–100.

[19] Nakamura A, Kaneko N, Villemagne VL, Kato T, Doecke J, Dore V, et al. High performance plasma amyloid-beta biomarkers for Alzheimer’s disease. Nature. 2018;554:249–54.

[20] Palmqvist S, Janelidze S, Stomrud E, Zetterberg H, Karl J, Zink K, et al. Performance of Fully Automated Plasma Assays as Screening Tests for Alzheimer Disease-Related beta-Amyloid Status. JAMA neurology. 2019.

[21] Palmqvist S, Insel PS, Stomrud E, Janelidze S, Zetterberg H, Brix B, et al. Cerebrospinal fluid and plasma biomarker trajectories with increasing amyloid deposition in Alzheimer’s disease. EMBO molecular medicine. 2019:e11170.

[22] van Oijen M, Hofman A, Soares HD, Koudstaal PJ, Breteler MM. Plasma Abeta(1-40) and Abeta(1-42) and the risk of dementia: a prospective case-cohort study. The Lancet Neurology. 2006;5:655–60.

[23] Lambert JC, Schraen-Maschke S, Richard F, Fievet N, Rouaud O, Berr C, et al. Association of plasma amyloid beta with risk of dementia: the prospective Three-City Study. Neurology. 2009;73:847–53.

[24] Shah NS, Vidal JS, Masaki K, Petrovitch H, Ross GW, Tilley C, et al. Midlife blood pressure, plasma beta-amyloid, and the risk for Alzheimer disease: the Honolulu Asia Aging Study. Hypertension. 2012;59:780–6.

[25] Chouraki V, Beiser A, Younkin L, Preis SR, Weinstein G, Hansson O, et al. Plasma amyloid-beta and risk of Alzheimer’s disease in the Framingham Heart Study. Alzheimer’s & dementia: the journal of the Alzheimer’s Association. 2015;11:249–57 e1.

[26] Toledo JB, Vanderstichele H, Figurski M, Aisen PS, Petersen RC, Weiner MW, et al. Factors affecting Abeta plasma levels and their utility as biomarkers in ADNI. Acta neuropathologica. 2011;122:401–13.

[27] Roberts KF, Elbert DL, Kasten TP, Patterson BW, Sigurdson WC, Connors RE, et al. Amyloid-beta efflux from the central nervous system into the plasma. Annals of neurology. 2014;76:837–44.

[28] Chouraki V, De Bruijn RF, Chapuis J, Bis JC, Reitz C, Schraen S, et al. A genome-wide association meta-analysis of plasma Abeta peptides concentrations in the elderly. Molecular psychiatry. 2014;19:1326–35.

[29] Figurski MJ, Waligorska T, Toledo J, Vanderstichele H, Korecka M, Lee VM, et al. Improved protocol for measurement of plasma beta-amyloid in longitudinal evaluation of Alzheimer’s Disease Neuroimaging Initiative study patients. Alzheimer’s & dementia: the journal of the Alzheimer’s Association. 2012;8:250–60.

[30] Ibrahim-Verbaas CA, Zorkoltseva IV, Amin N, Schuur M, Coppus AM, Isaacs A, et al. Linkage analysis for plasma amyloid beta levels in persons with hypertension implicates Abeta-40 levels to presenilin 2. Human genetics. 2012;131:1869–76.

[31] Reitz C, Cheng R, Schupf N, Lee JH, Mehta PD, Rogaeva E, et al. Association between variants in IDE-KIF11-HHEX and plasma amyloid beta levels. Neurobiology of aging. 2012;33:199 e13–7.

[32] Winkler TW, Day FR, Croteau-Chonka DC, Wood AR, Locke AE, Magi R, et al. Quality control and conduct of genome-wide association meta-analyses. Nature protocols. 2014;9:1192–212.

[33] Willer CJ, Li Y, Abecasis GR. METAL: fast and efficient meta-analysis of genomewide association scans. Bioinformatics. 2010;26:2190–1.

[34] de Leeuw CA, Mooij JM, Heskes T, Posthuma D. MAGMA: generalized gene-set analysis of GWAS data. PLoS computational biology. 2015;11:e1004219.

[35] Becker JA, Hedden T, Carmasin J, Maye J, Rentz DM, Putcha D, et al. Amyloid-beta associated cortical thinning in clinically normal elderly. Ann Neurol. 2011;69:1032–42.

[36] Splansky GL, Corey D, Yang Q, Atwood LD, Cupples LA, Benjamin EJ, et al. The Third Generation Cohort of the National Heart, Lung, and Blood Institute’s Framingham Heart Study: design, recruitment, and initial examination. American journal of epidemiology. 2007;165:1328–35.

[37] de Rojas I, Moreno-Grau S, Tesi N B. G-B, Andrade V, Jansen I, et al. Common variants in Alzheimer’s disease: Novel association of six genetic variants with AD and risk stratification by polygenic risk scores. medRxiv. 2019.

[38] Yamazaki Y, Zhao N, Caulfield TR, Liu CC, Bu G. Apolipoprotein E and Alzheimer disease: pathobiology and targeting strategies. Nature reviews Neurology. 2019;15:501–18.

[39] Grothe MJ, Villeneuve S, Dyrba M, Bartres-Faz D, Wirth M, Alzheimer’s Disease Neuroimaging I. Multimodal characterization of older APOE2 carriers reveals selective reduction of amyloid load. Neurology. 2017;88:569–76.

[40] Zhao N, Liu CC, Qiao W, Bu G. Apolipoprotein E, Receptors, and Modulation of Alzheimer’s Disease. Biological psychiatry. 2018;83:347–57.

[41] Miller KK, Hoffer A, Svoboda KR, Lupica CR. Cholecystokinin increases GABA release by inhibiting a resting K+ conductance in hippocampal interneurons. The Journal of neuroscience: the official journal of the Society for Neuroscience. 1997;17:4994–5003.

[42] Li QS, Parrado AR, Samtani MN, Narayan VA, Alzheimer’s Disease Neuroimaging I. Variations in the FRA10AC1 Fragile Site and 15q21 Are Associated with Cerebrospinal Fluid Abeta1-42 Level. PloS one. 2015;10:e0134000.

[43] Hnisz D, Abraham BJ, Lee TI, Lau A, Saint-Andre V, Sigova AA, et al. Super-enhancers in the control of cell identity and disease. Cell. 2013;155:934–47.

[44] Hampel H, O’Bryant SE, Molinuevo JL, Zetterberg H, Masters CL, Lista S, et al. Blood-based biomarkers for Alzheimer disease: mapping the road to the clinic. Nature reviews Neurology. 2018;14:639–52.

[45] Olsson B, Lautner R, Andreasson U, Ohrfelt A, Portelius E, Bjerke M, et al. CSF and blood biomarkers for the diagnosis of Alzheimer’s disease: a systematic review and meta-analysis. The Lancet Neurology. 2016;15:673–84.

[46] Sakai K, Senda T, Hata R, Kuroda M, Hasegawa M, Kato M, et al. Patients that have Undergone Hemodialysis Exhibit Lower Amyloid Deposition in the Brain: Evidence Supporting a Therapeutic Strategy for Alzheimer’s Disease by Removal of Blood Amyloid. Journal of Alzheimer’s disease: JAD. 2016;51:997–1002.

[47] Jin WS, Shen LL, Bu XL, Zhang WW, Chen SH, Huang ZL, et al. Peritoneal dialysis reduces amyloid-beta plasma levels in humans and attenuates Alzheimer-associated phenotypes in an APP/PS1 mouse model. Acta neuropathologica. 2017;134:207–20.

[48] Mullard A. BACE inhibitor bust in Alzheimer trial. Nature reviews Drug discovery. 2017;16:155.

[49] Fullwood NJ, Hayashi Y, Allsop D. Plasma amyloid-beta concentrations in Alzheimer’s disease: an alternative hypothesis. The Lancet Neurology. 2006;5:1000–1; author reply 2-3.

