## Supplementary Materials and Methods for "Plasma amyloid β levels are driven by genetic variants near *APOE, BACE1, APP, PSEN2:* A genome-wide association study in over 12,000 non-demented participants"

**1. Study Populations**

1.1. Framingham Heart Study

The Framingham Heart Study (FHS) is an ongoing community-based prospective cohort study of cardiovascular disease and its risk factors. It was initiated in 1948 with the enrollment of 5209 women and men aged 28 to 74 years (Original Cohort) [1]. Original cohort participants are reassessed biennially at a comprehensive core examination and have been examined 32 times to date [2]. In 1971, offspring of the original cohort and the spouses of these offspring (n = 5124, age 5-70 years, 3548 biological offspring, 1576 offspring spouses) were enrolled in the Framingham Offspring Cohort [3]. They have been examined every 4 to 8 years since, 9 times to date, for a core examination [4]. The Framingham Gen 3 cohort participants (n=4095, enrolled in 2002) are grandchildren of the Original cohort and children of the Offspring cohort [5]. All cohorts have been under ongoing surveillance for cognitive decline and dementia since 1975.

A total of 4,039 participants who attended the 23rd Original cohort examination (1992–1996, n=772) or the 7th Offspring examination (1998–2001, n=3267) had plasma Aβ1-42 and Aβ1-40 measured. We excluded participants with missing data for dementia (n=55) and genetic data (n=442), participants with extreme plasma Aβ values (n=3) and participants with prevalent dementia (n=42), yielding a subsample of 3523 participants for genome-wide association study of plasma Aβ concentrations.

Plasma Aβ levels were also measured in the 3^rd^ Generation Study as a separate effort. A total of 3401 participants who attended the 2^nd^ Gen3 examination (2008-2011, n=3411) had plasma Aβ1-42 and Aβ1-40 measured. We excluded participants with missing data for genetic data (n=188), participants with extreme plasma Aβ values (n=1) and participants with prevalent dementia (n=0), yielding a subsample of 3212 participants for genomewide association study of plasma Aβ concentrations.

The study protocol was approved by the Institutional Review Board of the Boston University Medical Center and all participants provided written informed consent.

1.2. Rotterdam Study

The Rotterdam Study is an ongoing, prospective, population-based cohort study investigating risk factors and incidence of cardiovascular, neurodegenerative, locomotor and ophthalmological diseases in elderly people [6]. From 1990–93, all 10 275 residents of Ommoord (a district of Rotterdam) aged 55 years or older were invited to participate in an extensive home interview and two visits to the research centre; 7 983 (78%) agreed. At the baseline clinical examination, blood samples were drawn from 7 050 individuals, of whom 7 047 underwent screening for dementia. Prevalent dementia was diagnosed in 334 of the latter. Hence, the cohort at risk of dementia comprised 6 713 participants. A random sub-cohort of 1 756 people was drawn from this source population for plasma Aβ concentration assessment [7]. After excluding those with missing genotype data, missing covariates, or outlying Aβ measures, 1549 individuals were included for this study. In addition, plasma Aβ levels were measured from blood drawn at the second visit to the research center in a sample of 491 participants as part of the NESTOR programme. After excluding those with missing genotypes, missing covariates, outlying Aβ measures and those already measured in the first batch, a total of 339 were included. Finally, plasma Aβ levels were measured during the third follow-up in all participants who had an earlier Aβ measure. An additional sample of 70 individuals that had previously been excluded now had complete measurements and was included as third batch. In total 1958 individuals from the Rotterdam study were included for this study. All participants gave informed consent. The study was approved of by the Erasmus University Medical Center Medical Ethics Committee.

1.3. Three City Study

The Three-City (3C) Study is a prospective cohort study of vascular risk factors for dementia. The methodology of the study has been described in detail elsewhere [8]. The 3C study's protocol was approved by the independent ethics committee at Kremlin-Bicêtre University Hospital (Paris). In 1999-2000, a sample of 9294 community dwellers aged 65 and over was selected from the electoral rolls of three French cities: Bordeaux (n=2104), Dijon (n=4931) and Montpellier (n=2259). Of these, 880 participants were excluded due to lack of a blood sample or lack of participation in any of the follow-up examinations. This left a sample of 8414 participants. For the present study, we focused on participants that were randomly selected as part of an ancillary case-cohort study (3C1, n=897) [9, 10], as well as two other consecutive random samples in which plasma Aβ levels were measured (3C2, n=1326 and 3C3, n=491). After excluding duplicates (n=224), participants with missing data for plasma Aβ (n=53) or genetic data (n=319), related individuals (n=88), participants with extreme values of plasma Aβ (n=12) or that did not pass genetic quality control (n=35) and participants with prevalent dementia (n=40), A total sample of 1954 participants (3C1, n=581, 3C2, n=1003, 3C3, n=370) was available for analysis.

1.4. Atherosclerosis Risk in Communities Study

ARIC was initiated in 1987 as a population-based cohort study of 15,792 middle-aged (45-64 years) participants drawn from four US communities (Washington County, MD; Forsyth County, NC; Jackson, MS; and suburban Minneapolis, MN) [11]. Four study visits were completed by 1999, with a fifth visit (ARIC-NCS; N=6,538) conducted in 2011-2013 . Plasma aβ, the phenotype for this investigation, was quantified on a subset (N=2,588) of ARIC-NCS enriched for cognitive impairment. All individuals exhibiting impaired cognitive status (defined as low mini-mental status exam score or low standardized score on any of five cognitive domains accompanied by cognitive decline on longitudinally administered tests) during the fifth exam, all participants with a brain MRI from a prior ARIC exam, and an age-stratified (<80 years, ≥80 years) random sample of the remaining cognitively normal participants from each field center were invited for plasma Aβ assessment. Plasma Aβ was simultaneously measured from blood samples obtained at the third (1993-1995) and fifth visits. This investigation focused on the fifth visit amyloid-β measurements from 830 cognitively normal participants of European ancestry who had genome-wide association data. An algorithmic syndromic diagnosis was used to define normal cognition. This algorithm synthesized information from the following assessments: pro-rated mini-mental status exam; longitudinal declines on the delayed word recall (DWRT), digit symbol substitution (DSST), and word fluency test (WFT: FAS); age-, race-, and education-adjusted z-scores on five cognitive domains (memory, executive, visuospatial, language, and attention); the Standard Clinical Dementia Rating Summary (CDRsb); and the functional activities questionnaire (FAQ). This algorithmic diagnosis yielded 1002 cognitively normal individuals of European ancestry who had blood samples sent for amyloid-β assessment. However, the final sample size (N=830) excluded individuals for genetic non-consent (N=2), prescriptions for anti-dementia drugs (N=3), lack of genome-wide association data (N=149), relatedness between participants (N=4), missing amyloid values (empty tubes or concentrations below the minimal detection threshold ; N=8), outlier amyloid values (>4 standard deviations from the mean; N=5), and inability to correct for plate effects (lone person on a plate; N=1).The ARIC study has been approved by the Institutional Review Board at each field center, namely Wake Forest Baptist Medical Center (Forsyth County, NC), University of Mississippi Medical Center (Jackson, MS), University of Minnesota (suburban Minneapolis, MN), and Johns Hopkins University (Washington County, MD). Participants provided written informed consent prior to each examination.

1.5. Epidemiological Prevention study Zoetermeer

The Zoetermeer Study, also called EPOZ (Epidemiologisch Preventief Onderzoek Zoetermeer,  translated: Epidemiological Preventive Study Zoetermeer), is a population-based prospective cohort study among 10,361 persons aged 5 to 91 years at the baseline year (1975) that was originally concerned with the prevalence of various chronic diseases [12]. All participants were included between 1975 and 1978.18 Zoetermeer is a suburban residential community (at that time, with about 55,000 inhabitants) that is near The Hague in the Netherlands. All participants gave informed consent. The study was approved of by the Erasmus University Medical Center Medical Ethics Committee.

For the current focus cohort, 909 subjects aged between 60 and 90 years who were randomly selected in strata of age (5-year strata), sex, and study from a larger pool of subjects in the appropriate age groups from the 2 primary studies. On agreement of participation, a list of contraindications (dementia, contraindications for magnetic resonance imaging [MRI] scanning, blindness) was reviewed to assess eligibility. Of these 909 individuals, 819 subjects were eligible. Among the eligible participants, 514 (63%) agreed to have an MRI brain scan as well as cognitive testing and each individual received up to three scans during 1995-2009 [13].

1.6. Washington Heights-Inwood Community Aging Project

The WHICAP study is a prospective, population-based study of aging and dementia in Medicare recipients aged 65 years and older residing in northern Manhattan (Washington Heights, Hamilton Heights, and Inwood). The study has been described in details elsewhere [14]. The cohort comprises non-Hispanic White, African American, and Hispanic participants. For the present study, we restricted the analyses to non-Hispanic Whites only (N=193). WHICAP participants undergo neuropsychological testings, medical and neurological examination, and a series of questionnaires approximately every 18 months. Formal diagnosis is reviewed in a weekly consensus conference attended by a group of co-investigators with extensive clinical expertise in diagnoses of dementia and MCI. The Columbia University Institutional Review Board reviewed and approved this project. All individuals provided written informed consent.

1.7. Alzheimer's Disease Neuroimaging Initiative

Data used in the preparation of this article were obtained from the Alzheimer’s Disease Neuroimaging Initiative (ADNI) database (adni.loni.usc.edu). The ADNI was launched in 2003 as a public-private partnership, led by Principal Investigator Michael W. Weiner, MD. The primary goal of ADNI has been to test whether serial magnetic resonance imaging (MRI), positron emission tomography (PET), other biological markers, and clinical and neuropsychological assessment can be combined to measure the progression of mild cognitive impairment (MCI) and early Alzheimer’s disease (AD). For up-to-date information, see www.adni-info.org.

For the present study, we included participants from the “Cognitively Normal” group. After applying the same criteria than in the 3C study, 173 participants were included in the analyses.

This study was approved by institutional review boards of all participating institutions and written informed consent was obtained from all participants or authorized representatives.

1.8. Eramus Rucphen Family Study

The Erasmus Rucphen Family (ERF) study is a population-based study in a genetically isolated population. All approximately 3,000 participants in this study are living descendants of 22 couples who, at the end of the nineteenth century, had at least six children baptized in the community church. Extensive genealogy data are available from the year 1600 AD. For this study, hypertensive subjects aged 55–75 years who did not have a history of stroke or dementia were selected from the study population. Of the 261 eligible individuals invited for this study, 135 agreed to participate [15]. All participants gave informed consent. The study was approved of by the Erasmus University Medical Center Medical Ethics Committee. Of the 135, 129 participants also had genomic data available and were included in the analysis.

**2. Plasma Aβ assessment**

2.1. Framingham Heart Study

EDTA plasma specimens used for the Aβ analyses were drawn into K3-EDTA evacuated specimen tubes, in the early afternoon in a supine nonfasting state for Original cohort, and in the morning in a supine fasting state for Offspring cohort. Specimen tubes were centrifuged for 30 minutes at 1850g at 4 degrees Celsius. Plasma was then separated from cells after centrifugation and placed at -80 degrees Celsius, within 90 minutes of venipuncture. The original aliquots consisted of 2mL of plasma in 3mL cryogenic storage vials for Original cohort samples and 700 µL of plasma in 1mL cryogenic storage vials for Offspring cohort samples. All specimens were stored at -80 degrees Celsius until they were aliquoted in March 2012 to be frozen and shipped for the assay. Therefore, specimens were thawed once prior to Aβ measurement. New aliquots consisted of 150 µL of plasma in 0.5 mL cryogenic storage vials. All samples were analyzed at the Department of molecular pharmacology and experimental therapeutics of the Mayo Clinic, Jacksonville, FL, from June to August 2012. Quantification of Aβ isoforms in plasma was performed using INNO-BIA plasma Aβ forms assays (Innogenetics NV, Ghent, Belgium), which is a multiplex microsphere-based Luminex xMAP technique that allows simultaneous analysis of Aβ1–40 and Aβ1–42 [16]. Measurements were done in duplicate in a randomly selected sample representing 9% of all samples. Intra-assay coefficients of variations (CV) for Aβ1-40 and Aβ1-42 were 3.2% and 2.6% and inter-assay CVs were 10.5% and 7.6%, respectively. Analysis of 146 phantom samples showed intraclass correlation coefficients of 0.916 and 0.943 and CV of 4.8% and 3.5%, respectively.

In Gen 3, plasma Aβ concentrations were measured following the same protocol between January and June 2014. Measurements were done in duplicate in a randomly selected sample representing 9% of all samples. Intra-assay coefficients of variations (CV) for Aβ1-40 and Aβ1-42 were 3.9% and 2.3% and inter-assay CVs were 6.8% and 6.6%, respectively. Analysis of 186 phantom samples showed intraclass correlation coefficients of 0.733 and 0.936 and CV of 7.9% and 4.9%, respectively.

### 2.2. Rotterdam Study

In the Rotterdam study, non-fasting blood samples obtained at baseline were placed in Vacutainer tubes (Becton Dickenson, Franklin Lakes, NJ, USA) containing sodium citrate. These samples were put on ice immediately and centrifuged within 60 min. Aliquots of plasma were stored at -80°C. Plasma Aβ concentrations were determined using a double-antibody sandwich enzyme-linked immunosorbent assay (ELISA) method (Pfizer, New York, NY, USA) [7]. The detection ranges were 10-1000 pg/mL for Aβ1-40 and 5-100 pg/mL for Aβ1-42.

### 2.3. Three City Study

In the 3C study, fasting plasma samples were collected in tubes containing sodium EDTA as an anticoagulant. After centrifugation, plasma samples were divided into aliquots in polypropylene tubes, stored at -80°C and only thawed immediately before Aβ quantification. The plasma Aβ peptide assay was performed using an INNO-BIA kit (Innogenetics NV, Ghent, Belgium) based on a multiplex xMAP (Luminex, Austin, TX, USA) technique.

### 2.4. Atherosclerosis Risk in Communities Study

Amyloid quantification was performed by the Department of Molecular Pharmacology and Experimental Therapeutics at Mayo Clinic, Jacksonville, FL, from August to December 2014. The INNO-BIA assay (INNOGENETICS N.V, Ghent, Belgium) used 69 plates to measure Aβ42 and Aβ40 levels. Beads (xMAP microspheres; conjugate 1A) bound to Aβ40 and Aβ42 emitted fluorescence detected by the Luminex 200 IS Total system. A five-parameter logistic regression model related the fluorescence intensities of six standards to their known amyloid concentrations. The resultant model predicted the concentrations of Aβ40 and Aβ42 from the measured fluorescence intensities in the samples. Intensities outside the range of the standards could not be inferred. The minimal detectable levels for Aβ42 and Aβ40 were 12 pg/ml and 15 pg/mL, respectively.

### 2.5. Epidemiological Prevention study Zoetermeer

Nonfasting blood samples were collected into sodium citrate containing Vacutainer blood collection tubes. These samples were put on ice directly and centrifuged within 60 minutes. Thereafter, aliquots of the samples were stored at -80˚C. Plasma Aβ levels were assessed using a double-antibody sandwich enzyme-linked immunosorbent assay method. The detection limits for Aβ1–40 and Aβ1–42 were 10 to 1,000 pg/ml and 5 to 100 pg/ml, respectively.

### 2.6. Washington Heights-Inwood Community Aging Project

Participants were asked to provide a 10-mL venous non-fasting blood sample (K3 EDTA lavendertop tube). Plasma levels of Aβ42 and Aβ40 were measured using a combination of monoclonal antibody 6E10 (specific to an epitope present on 1–16 amino acid residues of Aβ) and rabbit antisera R165 (vs Aβ42) and R162 (vs Aβ40) in a double antibody sandwich ELISA. The detection limit for these assays was 5 pg/mL for Aβ40 and 10 pg/mL for Aβ42. Aβ40 and Aβ42 levels from each sample were measured twice using separate aliquots. Reliability between measurements was substantial for both peptides (r = 0.93 and r = 0.97 for Aβ40 and Aβ42, p = 0.001), and we used the mean of the two measurements in statistical analyses.

### 2.7. Alzheimer's Disease Neuroimaging Initiative

In the ADNI study, plasma samples were obtained from the ADNI biofluid repository at the University of Pennsylvania. The plasma samples were collected at the participating centers. After overnight fasting, plasma was collected in the morning by venipuncture and placed in Vacutainer tubes containing potassium K3 ethylene tetraacetate as an anticoagulant. After centrifugation, samples were placed in polypropylene transfer tubes (13 mL, Sarstedt Inc., Newton, NC, USA catalogue number 60.541), frozen and shipped on dry ice to the UPenn Biomarker Core Laboratory, where they were stored temporarily at -80°C. Within several weeks of receipt, the samples were thawed, aliquoted by 500 μL into polypropylene tubes (1.5 mL, Thermo Fisher Scientific, Waltham, MA, USA catalogue number 05-408-129) and stored at -80°C pending biochemical analyses. The plasma levels of Aβ1-40 and Aβ1-42 were quantified with the INNO-BIA kit (Innogenetics NV, Ghent, Belgium).

### 2.8. Eramus Rucphen Family Study

In the ERF study non-fasting blood samples were obtained in EDTA tubes and immediately cooled on ice. Plasma was extracted and stored at −80 °C. Plasma Aβ concentrations were measured with a fluorimetric bead-based immunoassay using xMAP® technology (Innogenetics®). Aβ40, Aβ42, and the truncated forms Aβn-42 and Aβn-40 were measured [15].

### 3. GWAS genotyping and 1000 Genomes Imputation

### 3.1. Framingham Heart Study

Genotyping was conducted by Affymetrix (Santa Clara, CA) using the Affymetrix 500K Mapping and Affymetrix 50K Supplemental Array. SNP data were filtered for a call rate ≥ 97%, a deviation from Hardy-Weinberg (HWE) ≥ 1x10^-6^, Mishap P ≥ 1x10^-9^, Mendel errors ≤ 100, and MAF ≥ 1%, prior to being imputed to the 1000 Genomes Phase I version 3 (August 2012) reference panel using MACH (version 1.0.16).

3.2. Rotterdam Study

The Rotterdam Study were genotyped using the Illumina 550K chip. The following exclusions were applied to identify a final set of SNPs that was used in this study: MAF < 0.05, SNP callrate < 0.95 and/or HWE p-value < 1×10-7. The QC was done per cohort. Imputation per cohort to the 1000 genomes phase I version 3 was performed with MaCH and Minimac.

3.3. Three City Study

3C study samples were genotyped using the Illumina Human610-Quad BeadChip. Prior to imputation, quality control was performed on SNVs and samples. These include the exclusion of samples with too much missingness (>5%), samples with a discordant sex between clinical and genetic information, samples with an outlying heterozygosity and samples of non european ancestry based on principal components analysis on Hapmap. SNVs with a low call rate (<98%), a Pvalue of Hardy-Weinberg < 1e-6, a Minor Allele Frequency < 1% were excluded. Pre-phasing was performed using Shapeit v2.r727 and imputation was performed on 1000 genomes phase I version 3 using Impute2 v2.3.0.

3.4. Atherosclerosis Risk in Communities Study

Self-reported whites were genotyped using the Affymetrix Genome-wide Human SNP Array 6.0. The genotyped data were checked for gender mismatch between clinically reported data and genetic markers, genetic outliers, failed Taqman concordance checks (if discordance between Taqman and Affymetrix genotypes were greater than 20%), and relatedness. Measured genotypes were pre-phased using the ShapeIt software (v1.r532 using parameters --states-phase 200). Imputation was conducted by IMPUTE2 using measured single nucleotide polymorphisms with minor allele frequencies ≥0.005, SNP missingness < 5%, and Hardy-Weinberg p-values >0.00001.After frequency and genotyping pruning, there were 682,749 autosomal SNPs used for imputations. IMPUTE 2 relied on the 1000 Genomes Phase I Integrated Release Version 3 reference panel, NCBI build 37 (hg19) positions, and chunks of size 5 Mb.

3.5. Epidemiological Prevention study Zoetermeer

Genotyping was performed in 450 individuals at the Erasmus Medical Center, using the Illumina Infinium Global Screening Array v1.0. Prior to imputation, QC was done with the exclusion of 1) low quality samples and variants with call rate <97.5%, 2) variants that were out of HWE (filter of 1 x 10-4), 3) samples for which evidence for excess of heterozygosity was present, 4) gender mismatch, 5) monomorphic SNPs and 6) variants with >20% difference of MAF between reference and population allele frequency. After QC, 422 sample remained among those, 397 individuals were available for the present analysis after exclusion. Imputation to the 1000 genomes phase I version 3 was performed using the Michigan Imputation Server.

3.6. Washington Heights-Inwood Community Aging Project

Non-Hispanic Whites, part of the WHICAP study, were genotyped using Illumina 660 chip. Prior to imputation, quality control was performed, excluding 1) related and duplicate sample (PLINK pi-hat >0.2); 2) samples with high missingness (>5%); 3) samples with a discordant sex between clinical and genetic information; 4) samples with an outlying heterozygosity. SNPs with a low call rate (<98%), a Hardy-Weinberg p-value < 1 x 10-6, a MAF < 1% were also excluded. Pre-phasing was performed using SHAPEIT and imputation was performed using 1000 genomes – phase I version 3 as reference panel and Impute2 v2.3.0 for imputation.

3.7. Alzheimer's Disease Neuroimaging Initiative

ADNI study samples were genotyped using the Illumina Human610-Quad BeadChip. Prior to imputation, quality control was performed on SNVs and samples. These include the exclusion of samples with too much missingness (>5%), samples with a discordant sex between clinical and genetic information, samples with an outlying heterozygosity and samples of non european ancestry based on principal components analysis on Hapmap and “non white” ethnicity as mentioned in the ADNI demographic database. SNVs with a low call rate (<98%), a Pvalue of Hardy-Weinberg < 1e-6, a Minor Allele Frequency < 1% were excluded. Pre-phasing was performed using Shapeit v2.r727 and imputation was performed on 1000 genomes phase I version 3 using Impute2 v2.3.0.

3.8. Erasmus Rucphen Family Study

In ERF genotyping was done on various Illumina and Affymetrix chips. QC was done separately for each chip. For most chips, the following QC criteria were applied: callrate > 0.98, per individual callrate > 0.96, Hardy-Weinberg equilibrium (HWE) p-value > 5×10-8 and minor allele frequency (MAF) > 0.005. IBS checks, sex chromosome checks and ethnicity checks were also performed. The imputation to the Genome of the Netherlands reference panel, release 4 was performed with MaCH (Mach 1.0.18.c) and minimac (2012.8.15).

**4. APOEε genotyping**

4.1. Framingham Heart Study

Leucocyte DNA was extracted from 5–10 ml of whole blood. APOE genotype was performed as described by Hixson and Vernier. A 244-bp sequence of the APOE gene including the two polymorphic sites was amplified by polymerase chain reaction (PCR) in a DNA Thermal Cycler (PTC-100, MJ Research, Watertown, MA), using oligonucleotide primers F4 and F6. Each reaction mixture was heated at 94°C for 2 min and followed by 35 cycles of amplification (94°C for 40 s, 62°C for 30 s and 72°C for 1 min). The PCR products were digested with 5 U Hha I and the fragments separated by electrophoresis on an 8% polyacrylamide nondenaturing gel. After electrophoresis the gel was treated with ethidium bromide for 30 min and DNA fragments were visualized by UV illumination.

4.2.Rotterdam Study

Genotyping for *APOE* was performed on coded DNA specimens without knowledge of the diagnosis as previously described [17]. Briefly, a polymerase chain reaction was conducted, and the amplification products were digested with HhaI. The resulting restriction fragments were separated using precast ExcelGel gels (Pharmacia Biotech, Uppsala, Sweden) and visualized by silver staining [17].

4.3. Three City Study

Blood was collected on EDTA K3 (Le Pont-De-Claix, France). DNA was extracted from white blood cells with the Puregene extraction kit (Gentra Systems, Inc., Minneapolis, MN). *APOE* genotyping was performed using the fluorogenic 5'-nuclease assay with TaqMan chemistry. The sequences of the primers and probe oligonucleotides were designed as previously described [18]. Amplification was performed in a final volume of 5 µL containing 20 ng/µL of DNA solution, 900 nM of each primer, 200 nM of each probe, and 2x TaqMan Universal PCR master mix (Applied Biosystems, Foster City, CA). In each assay, controls for the wild type and mutations were included. Reaction mixtures were loaded into 384-well plates and placed in a GeneAmp PCR system 9700 (Applied Biosystems). The PCR conditions were as follows: initial denaturation at 95 °C for 10 minutes followed by 48 cycles of denaturation (92 °C for 15 seconds), annealing, and extension in one step (60 °C for 60 seconds). After cycling, genotyping was carried out on the ABI Prism Sequence detection system 7900 (Applied Biosystems).

4.4. Atherosclerosis Risk in Communities Study

The apolipoprotein-E (APOE) genotypes were determined via TaqMan assays (Applied Biosystems, Foster City, CA). Oligonucleotide sequences for polymerase chain reaction primers and TaqMan probes are available by request. The standard TaqMan assay can detect a maximum of two single nucleotide polymorphisms per reaction, thus the APOE variants at codons 112 (rs429358) and 158 (rs7412) were determined separately. The single nucleotide polymorphisms at the two positions were combined to generate six APOE genotypes (ε2/ε2, ε2/ε3, ε2/ε4, ε3/ε3, ε3/ε4, ε4/ε4). The ABI 7700 Sequence Detection System (Applied Biosystems, Foster City, CA) was employed to detect alleles and call genotypes. After applying the exclusion criteria, 96.6% (802/830) of the participants had non-missing APOE genotypes.

4.5. Epidemiological Prevention study Zoetermeer

*APOE*ε genotypes were assessed by a polymerase chain reaction on coded genomic DNA samples using a bi-allelic Taqman assay (rs7412 and rs429358).

4.6. Washington Heights-Inwood Community Aging Project

APOE genotyping employed standard PCR–restriction fragment length polymorphism methods using HhaI (CfoI) digestion of an APOE genomic PCR product spanning the polymorphic (cys/arg) sites at codons 112 and 158 [19]. Acrylamide gel electrophoresis was used to assess and document the restriction fragment sizes.

4.7. Alzheimer's Disease Neuroimaging Initiative

*APOE* genotyping was performed at the time of participant enrollment. The two SNPs (rs429358, rs7412) that define the ε2, ε3, and ε4 alleles were genotyped at the National Cell Repository for Alzheimer’s Disease (NCRAD) using DNA extracted by Cogenics from a 3 mL aliquot of EDTA blood.

4.8. Eramus Rucphen Family Study

Subjects were genotyped for the *APOE* ε2/3/4 polymorphism with two TaqMan allelic discrimination Assays-By-Design (Applied Biosystems, Foster City,CA), targeting SNPs in the 112th (rs429358) and 158th (rs7412) amino acids of the ApoE gene as previously described [20].

**5. PET imaging**

Amyloid PET imaging was carried out using the 11C-Pittsburgh Compound B (PiB) tracer on 193 suitably consented FHS Generation 3 participants who had attended the 2^nd^ quadrennial examination. PET imaging was conducted at the Massachusetts General Hospital on a CTI/Siemens EXACT HR+ PET scanner. The protocol had been approved by the Institutional review boards of the Boston Medical Center and the Massachusetts General Hospital. Prior to amyloid imaging, participants completed computer tomography for attenuation correction and structural brain MRI (on a 3T Phillips Acheiva scanner) for co-registration. PiB PET images were acquired with a 10 to 15 mCi bolus injection followed by a 60-minute dynamic acquisition in 69 vol (12 × 15 s, 57 × 60 s). PET data were reconstructed, attenuation corrected, scatter corrected, and evaluated frame by frame for excessive head motion. Data from the first 8 min post injection was used to co-register the data to the T1 image for each subject. A summary distribution volume ratio (DVR) was computed with Logan plotting was derived from a target region composed of frontal, lateral and retrosplenial tracer uptake (FLR) and a cerebellar grey reference region. Persons with a DVR across these regions of >1.36 were categorized as having significant amyloid burden and a positive scan. APOE genotyping in all participants had been completed using a Taqman assay and we confirmed congruence between these assessments and the status identified at genome-wide genotyping with imputation.

**6. References**

[1] Dawber TR, Meadors GF, Moore FE, Jr. Epidemiological approaches to heart disease: the Framingham Study. American journal of public health and the nation's health. 1951;41:279-81.

[2] Farmer ME, White LR, Kittner SJ, Kaplan E, Moes E, McNamara P, et al. Neuropsychological test performance in Framingham: a descriptive study. Psychological reports. 1987;60:1023-40.

[3] Feinleib M, Kannel WB, Garrison RJ, McNamara PM, Castelli WP. The Framingham Offspring Study. Design and preliminary data. Preventive medicine. 1975;4:518-25.

[4] Au R, Seshadri S, Wolf PA, Elias M, Elias P, Sullivan L, et al. New norms for a new generation: cognitive performance in the framingham offspring cohort. Experimental aging research. 2004;30:333-58.

[5] Splansky GL, Corey D, Yang Q, Atwood LD, Cupples LA, Benjamin EJ, et al. The Third Generation Cohort of the National Heart, Lung, and Blood Institute's Framingham Heart Study: design, recruitment, and initial examination. American journal of epidemiology. 2007;165:1328-35.

[6] Hofman A, Brusselle GG, Darwish Murad S, van Duijn CM, Franco OH, Goedegebure A, et al. The Rotterdam Study: 2016 objectives and design update. European journal of epidemiology. 2015;30:661-708.

[7] van Oijen M, Hofman A, Soares HD, Koudstaal PJ, Breteler MM. Plasma Abeta(1-40) and Abeta(1-42) and the risk of dementia: a prospective case-cohort study. The Lancet Neurology. 2006;5:655-60.

[8] Group CS. Vascular factors and risk of dementia: design of the Three-City Study and baseline characteristics of the study population. Neuroepidemiology. 2003;22:316-25.

[9] Carcaillon L, Gaussem P, Ducimetiere P, Giroud M, Ritchie K, Dartigues JF, et al. Elevated plasma fibrin D-dimer as a risk factor for vascular dementia: the Three-City cohort study. Journal of thrombosis and haemostasis : JTH. 2009;7:1972-8.

[10] Lambert JC, Schraen-Maschke S, Richard F, Fievet N, Rouaud O, Berr C, et al. Association of plasma amyloid beta with risk of dementia: the prospective Three-City Study. Neurology. 2009;73:847-53.

[11] The Atherosclerosis Risk in Communities (ARIC) Study: design and objectives. The ARIC investigators. American journal of epidemiology. 1989;129:687-702.

[12] Hofman A, Boomsma F, Schalekamp MA, Valkenburg HA. Raised blood pressure and plasma noradrenaline concentrations in teenagers and young adults selected from an open population. British medical journal. 1979;1:1536-8.

[13] de Groot JC, de Leeuw FE, Oudkerk M, van Gijn J, Hofman A, Jolles J, et al. Cerebral white matter lesions and cognitive function: the Rotterdam Scan Study. Annals of neurology. 2000;47:145-51.

[14] Tang MX, Cross P, Andrews H, Jacobs DM, Small S, Bell K, et al. Incidence of AD in African-Americans, Caribbean Hispanics, and Caucasians in northern Manhattan. Neurology. 2001;56:49-56.

[15] Ibrahim-Verbaas CA, Zorkoltseva IV, Amin N, Schuur M, Coppus AM, Isaacs A, et al. Linkage analysis for plasma amyloid beta levels in persons with hypertension implicates Abeta-40 levels to presenilin 2. Human genetics. 2012;131:1869-76.

[16] Blennow K, De Meyer G, Hansson O, Minthon L, Wallin A, Zetterberg H, et al. Evolution of Abeta42 and Abeta40 levels and Abeta42/Abeta40 ratio in plasma during progression of Alzheimer's disease: a multicenter assessment. The journal of nutrition, health & aging. 2009;13:205-8.

[17] Slooter AJ, Cruts M, Kalmijn S, Hofman A, Breteler MM, Van Broeckhoven C, et al. Risk estimates of dementia by apolipoprotein E genotypes from a population-based incidence study: the Rotterdam Study. Archives of neurology. 1998;55:964-8.

[18] Koch W, Ehrenhaft A, Griesser K, Pfeufer A, Muller J, Schomig A, et al. TaqMan systems for genotyping of disease-related polymorphisms present in the gene encoding apolipoprotein E. Clinical chemistry and laboratory medicine. 2002;40:1123-31.

[19] Hixson JE, Vernier DT. Restriction isotyping of human apolipoprotein E by gene amplification and cleavage with HhaI. Journal of lipid research. 1990;31:545-8.

[20] Isaacs A, Sayed-Tabatabaei FA, Aulchenko YS, Zillikens MC, Sijbrands EJ, Schut AF, et al. Heritabilities, apolipoprotein E, and effects of inbreeding on plasma lipids in a genetically isolated population: the Erasmus Rucphen Family Study. European journal of epidemiology. 2007;22:99-105.
