## Supplementary Figures for "Plasma amyloid β levels are driven by genetic variants near *APOE, BACE1, APP, PSEN2:* A genome-wide association study in over 12,000 non-demented participants"

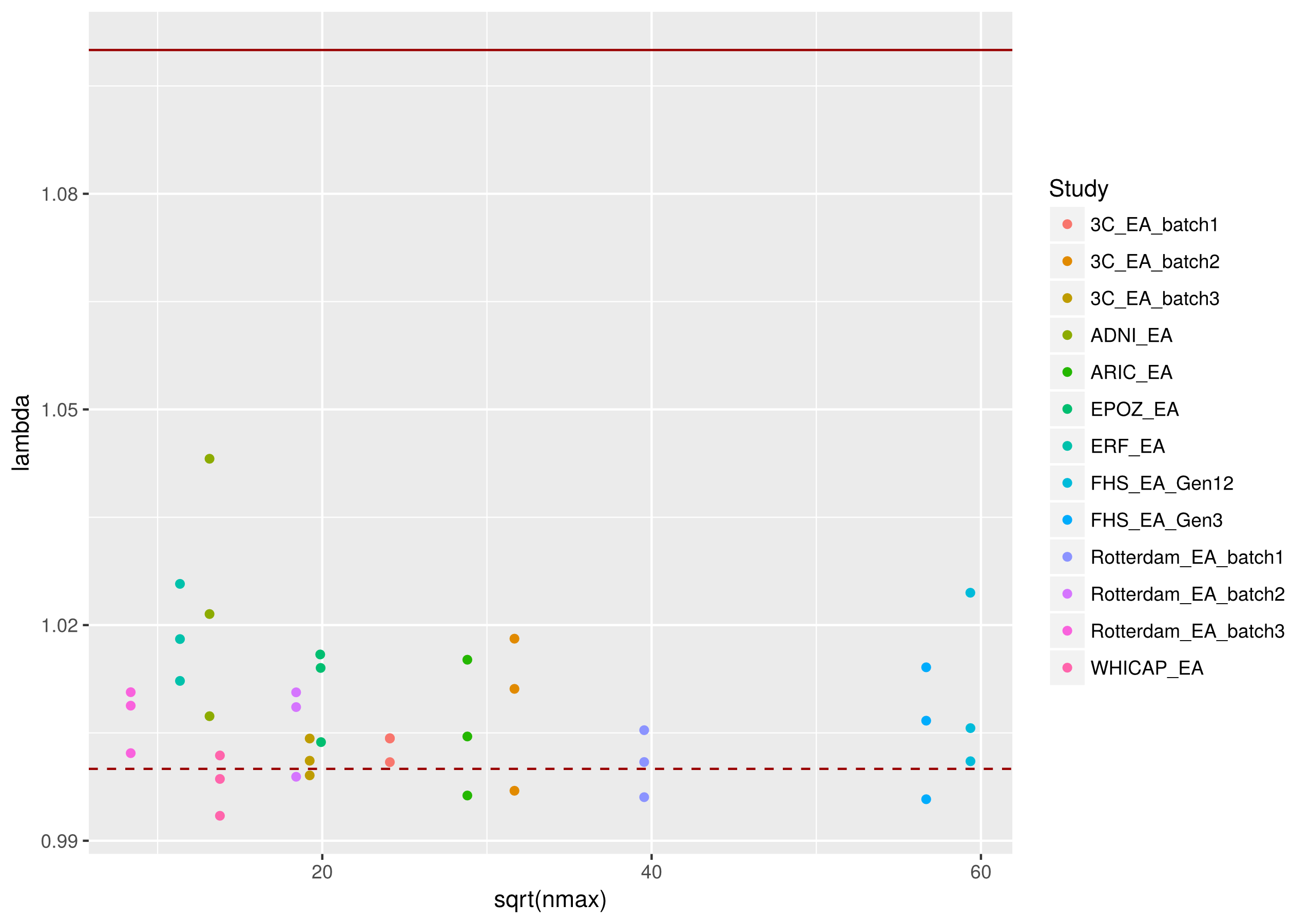
Supplementary Figure 1. Genomic inflation factors (λ) of individual GWAS of plasma Aβ1-40, Aβ1-42 and Aβ1-42/Aβ1-40 ratio according to sample size

Supplementary Figure 2: Verification of the homogeneity of plasma Aβ measures across studies after inverse-normal transformation

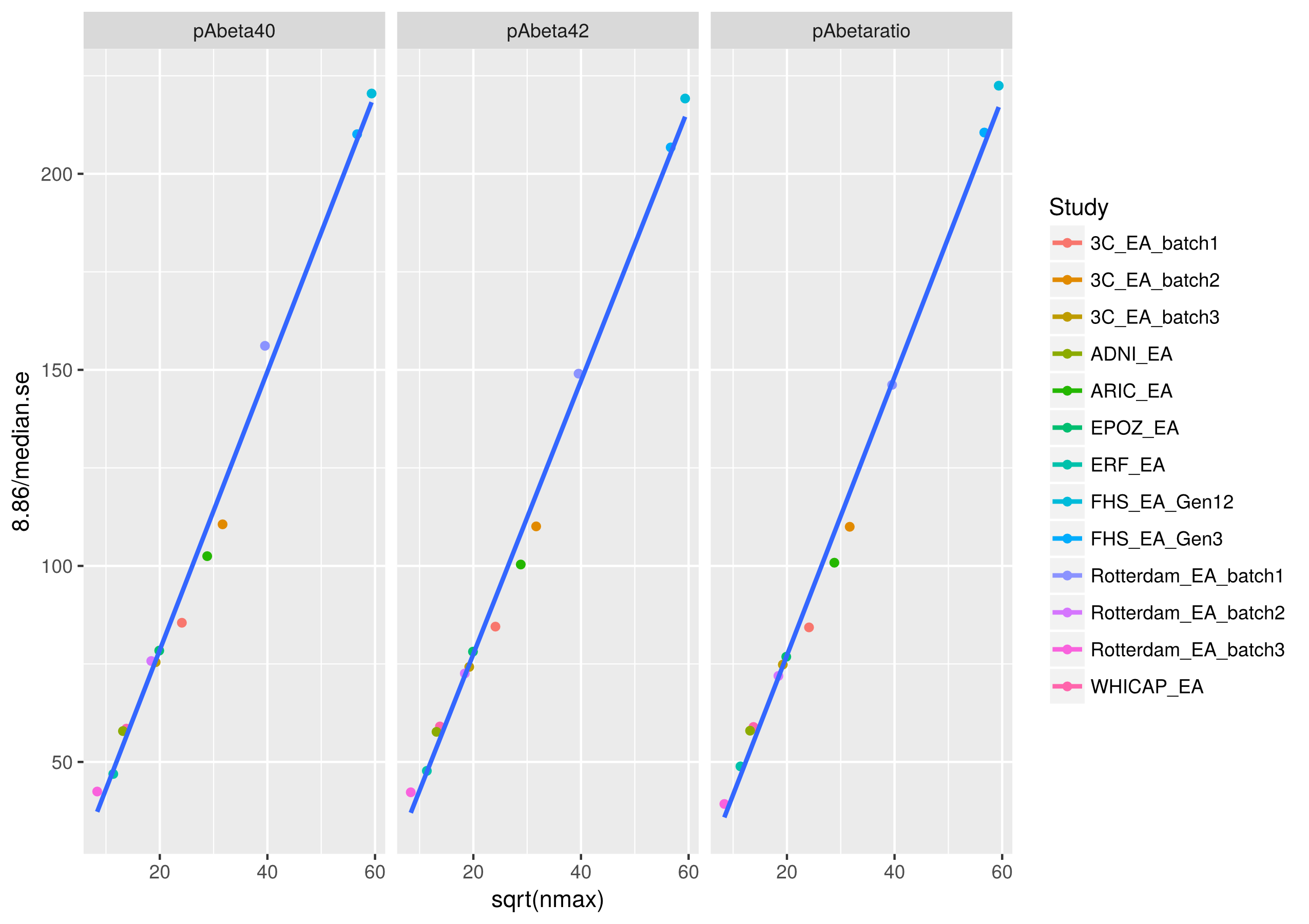

Supplementary Figure 3. Quantile-quantile plot of the genome-wide meta-analysis of plasma Aβ1-40 levels.

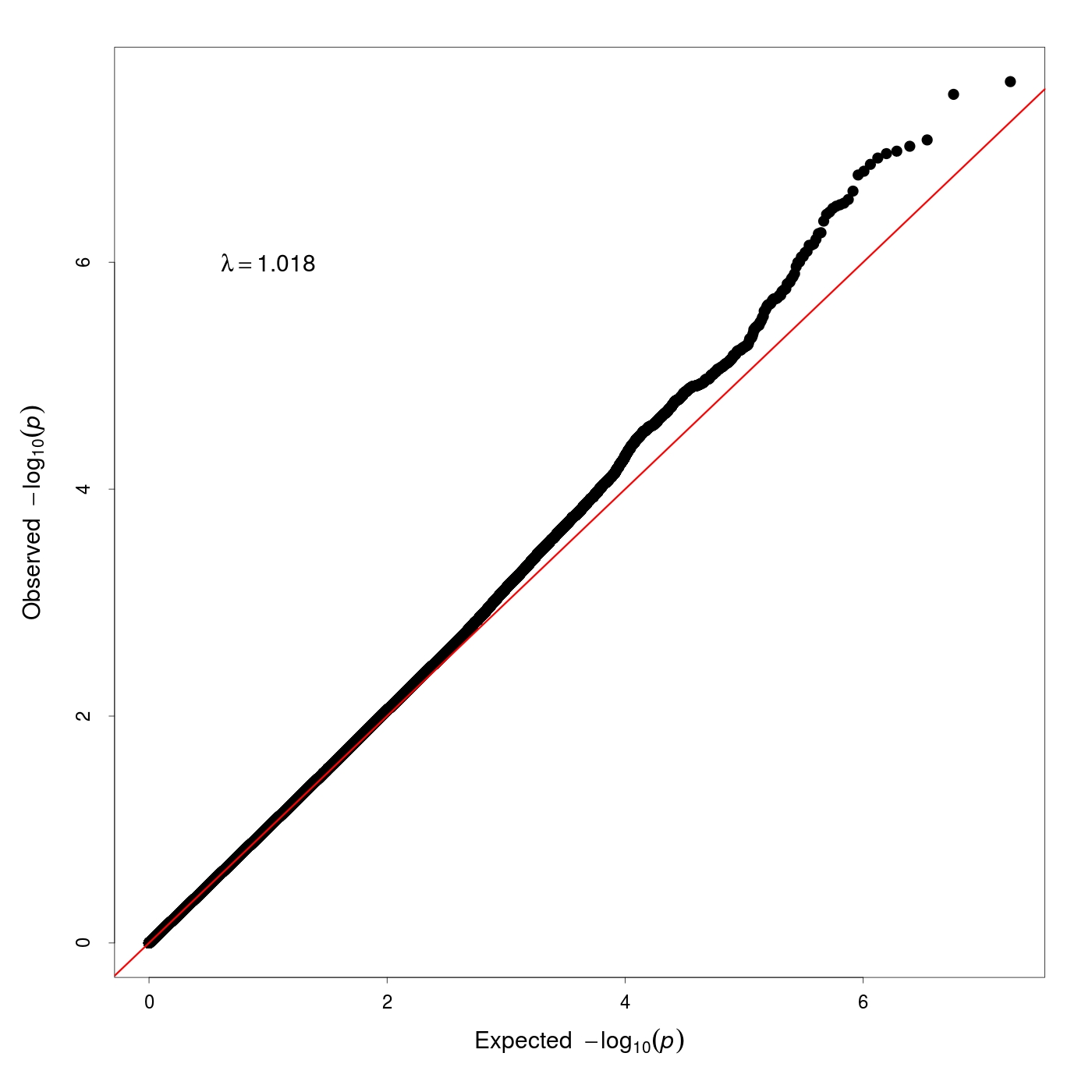

Supplementary Figure 4. Quantile-quantile plot of the genome-wide meta-analysis of plasma Aβ1-42 levels.

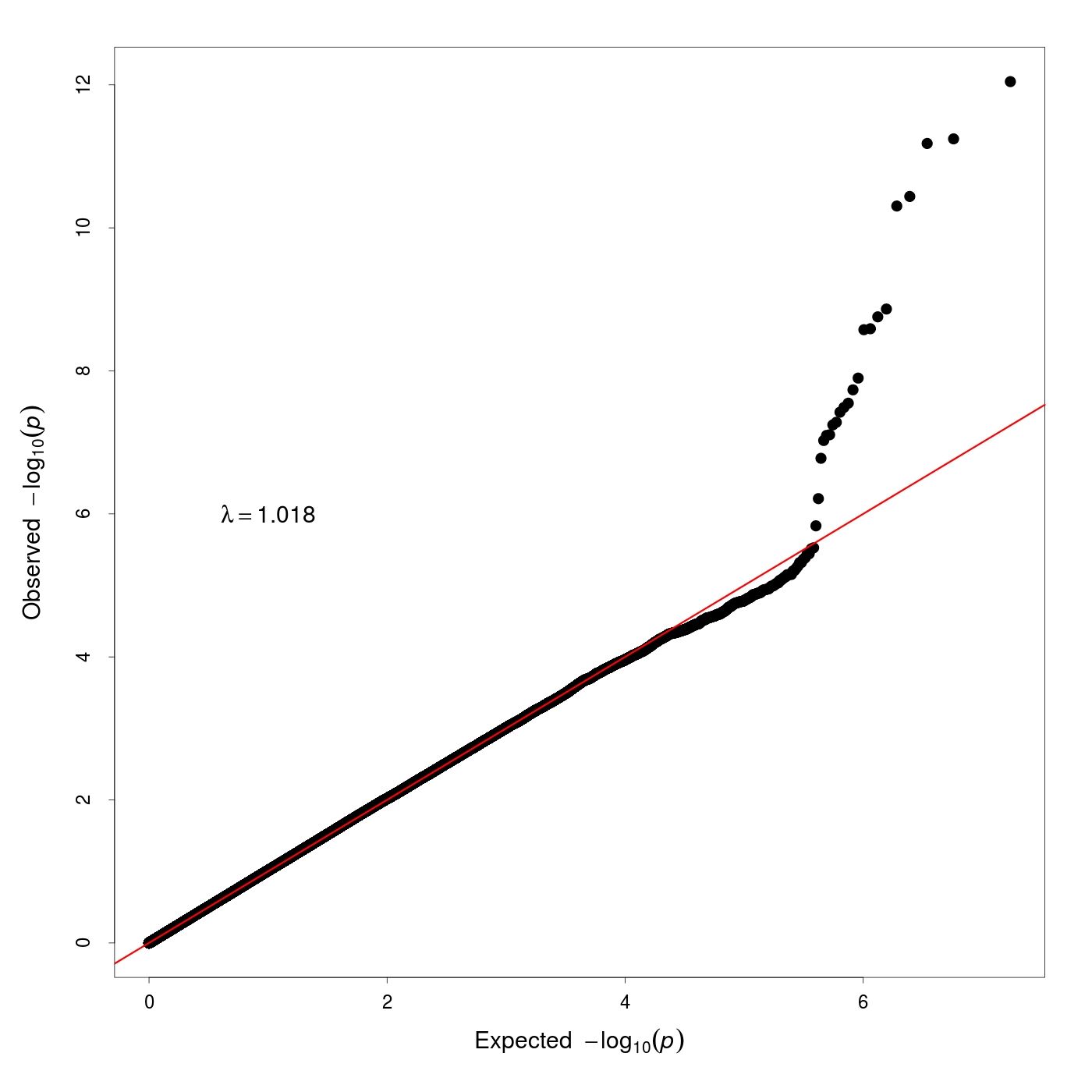

Supplementary Figure 5. Quantile-quantile plot of the genome-wide meta-analysis of plasma Aβ1-42/Aβ1-40 ratio.

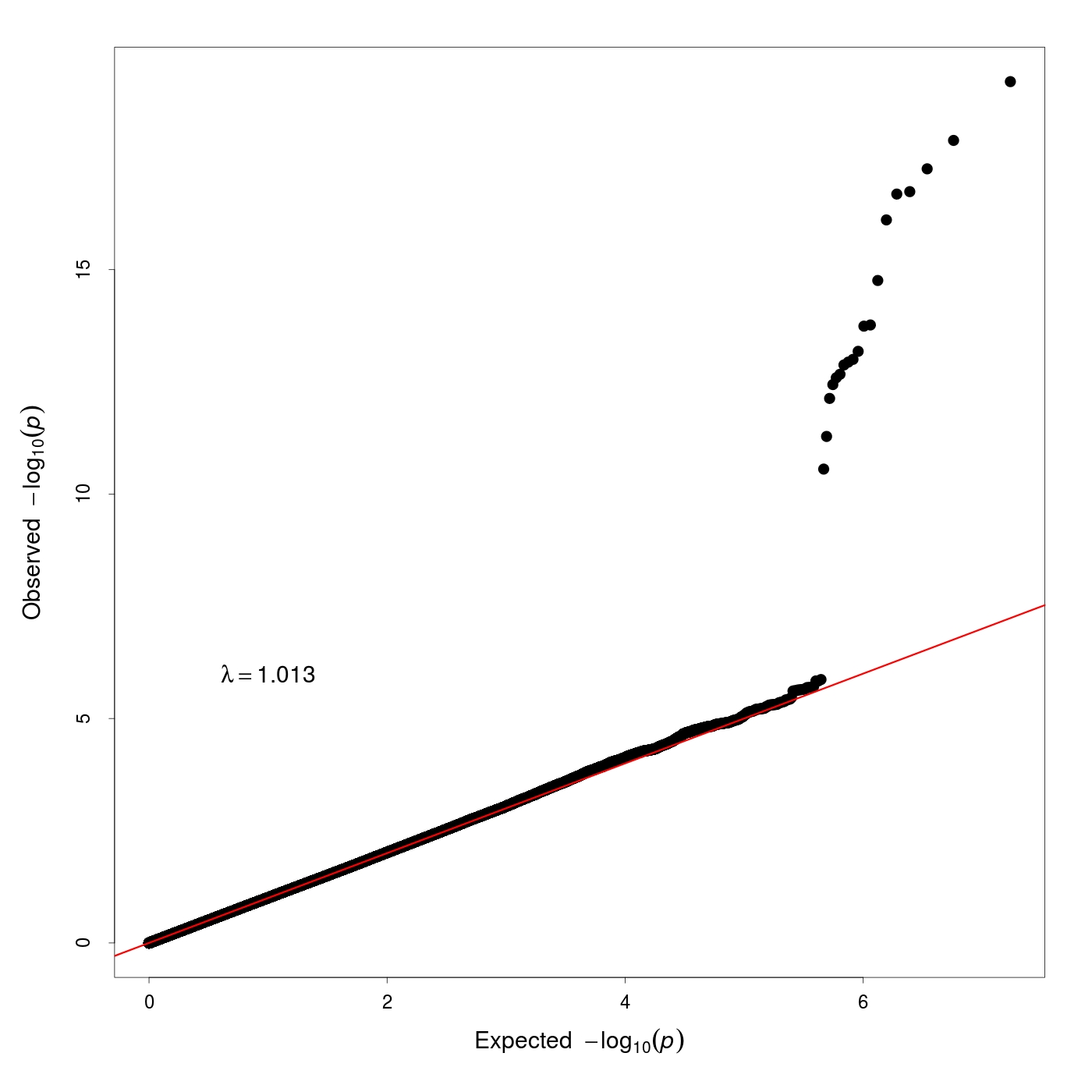

Supplementary Figure 6. Circos plot of the association results of plasma Aβ1-40 levels.

The first track shows the autosomal chromosomes. The second track (blue dots) indicates the –log_10_(p) of the SNP meta-analysis results. The red and dark green dotted lines represent the significative (p=5x10^-8^) and suggestive (p=1x10^-5^) thresholds for SNP association, respectively. The third track (brown dots) indicates the –log_10_(p) of the genes association results. The red dotted line represent the significative (p=2.76x10^-6^) threshold for gene association.

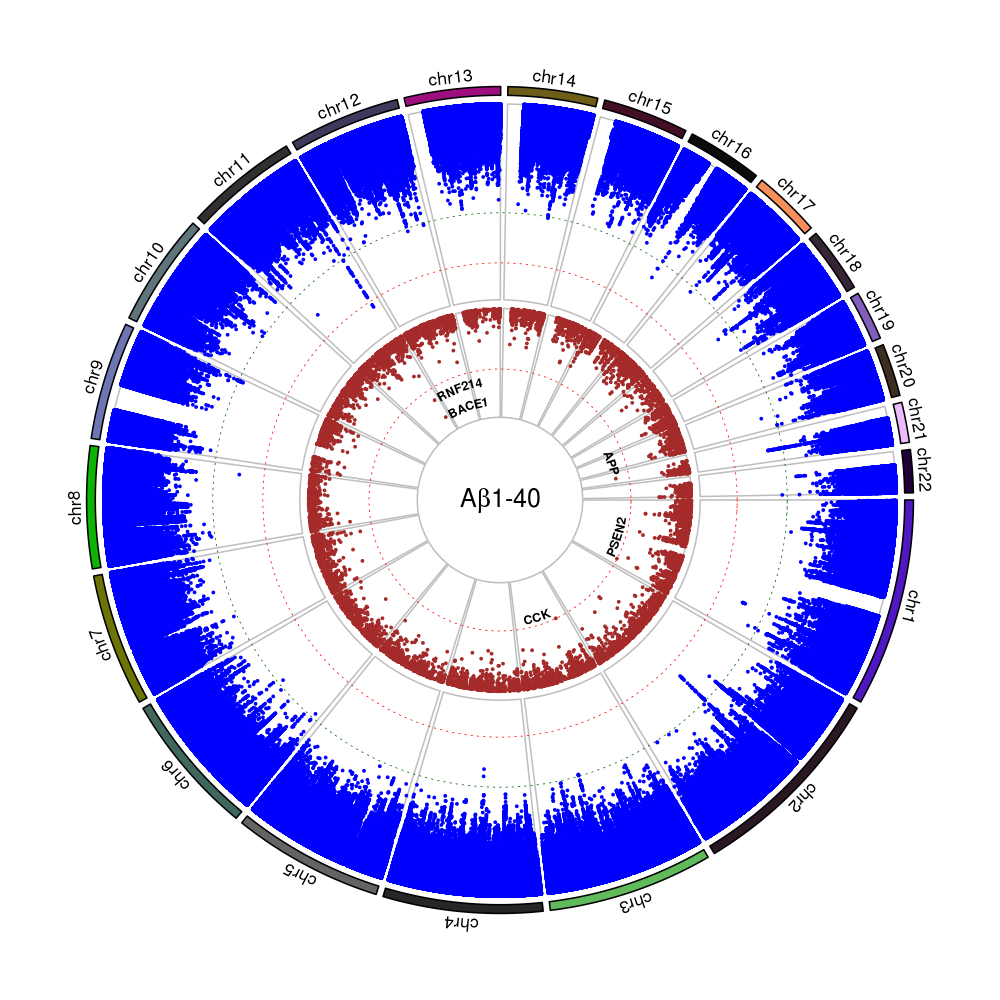

Supplementary Figure 7. Circos plot of the association results of plasma Aβ1-42 levels.

The first track shows the autosomal chromosomes. The second track (blue dots) indicates the –log_10_(p) of the SNP meta-analysis results. The red and dark green dotted lines represent the significative (p=5x10^-8^) and suggestive (p=1x10^-5^) thresholds for SNP association, respectively. The third track (brown dots) indicates the –log_10_(p) of the genes association results. The red dotted line represent the significative (p=2.76x10^-6^) threshold for gene association.

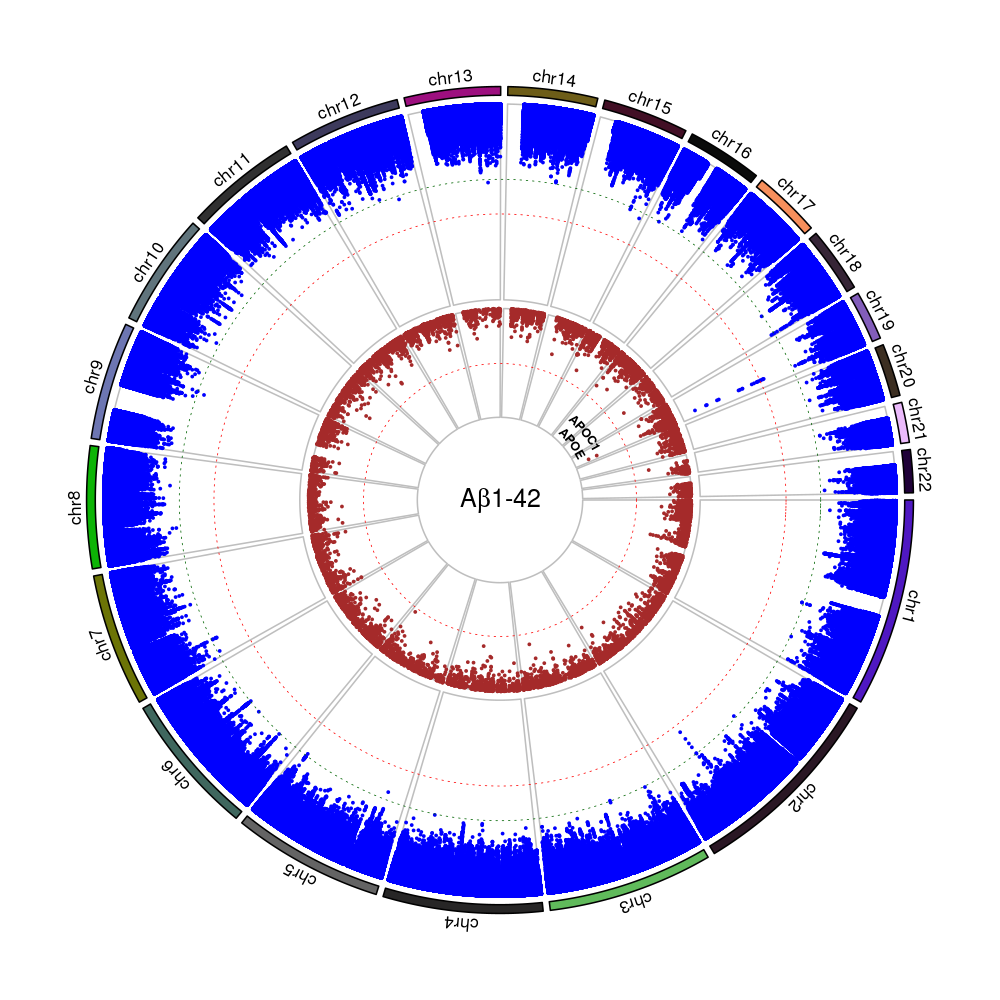

Supplementary Figure 8. Circos plot of the association results of plasma Aβ1-42/Aβ1-40 ratio.

The first track shows the autosomal chromosomes. The second track (blue dots) indicates the –log_10_(p) of the SNP meta-analysis results. The red and dark green dotted lines represent the significative (p=5x10^-8^) and suggestive (p=1x10^-5^) thresholds for SNP association, respectively. The third track (brown dots) indicates the –log_10_(p) of the genes association results. The red dotted line represent the significative (p=2.76x10^-6^) threshold for gene association.

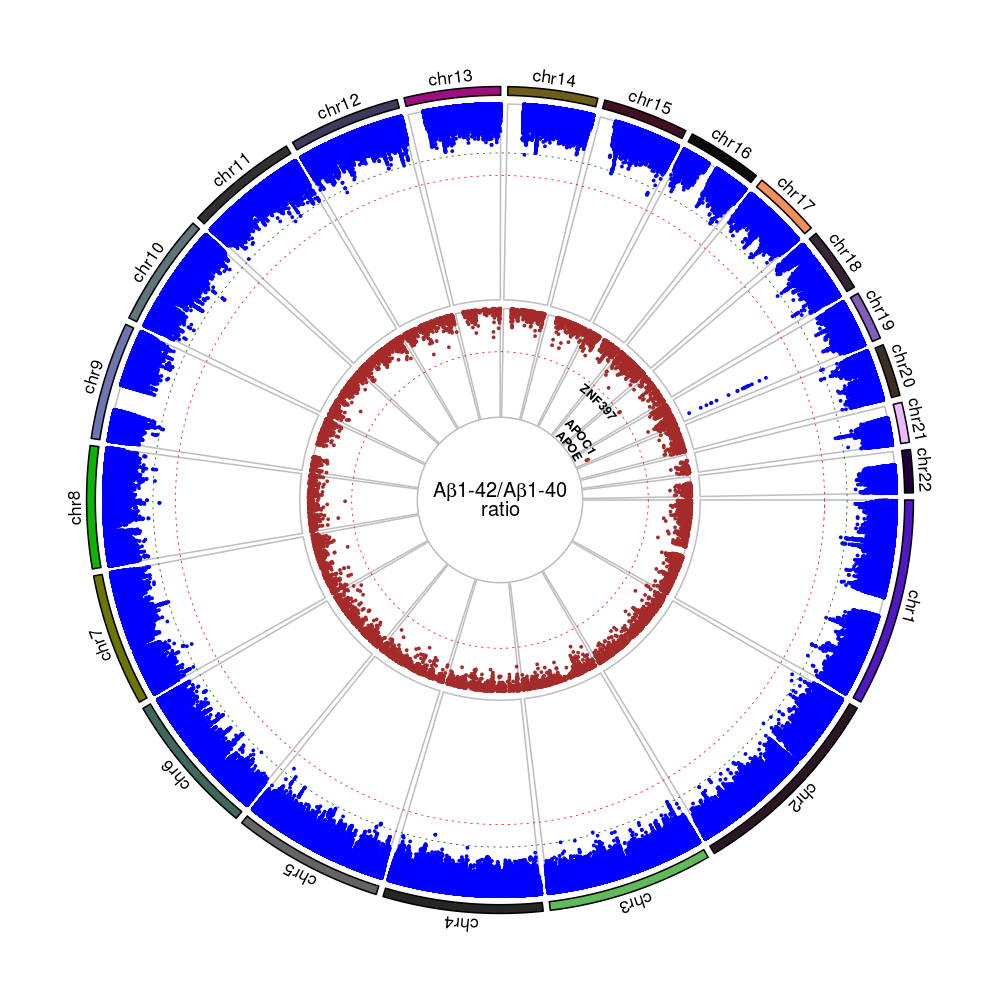

Supplementary Figure 9. Association of top hits with plasma Aβ levels

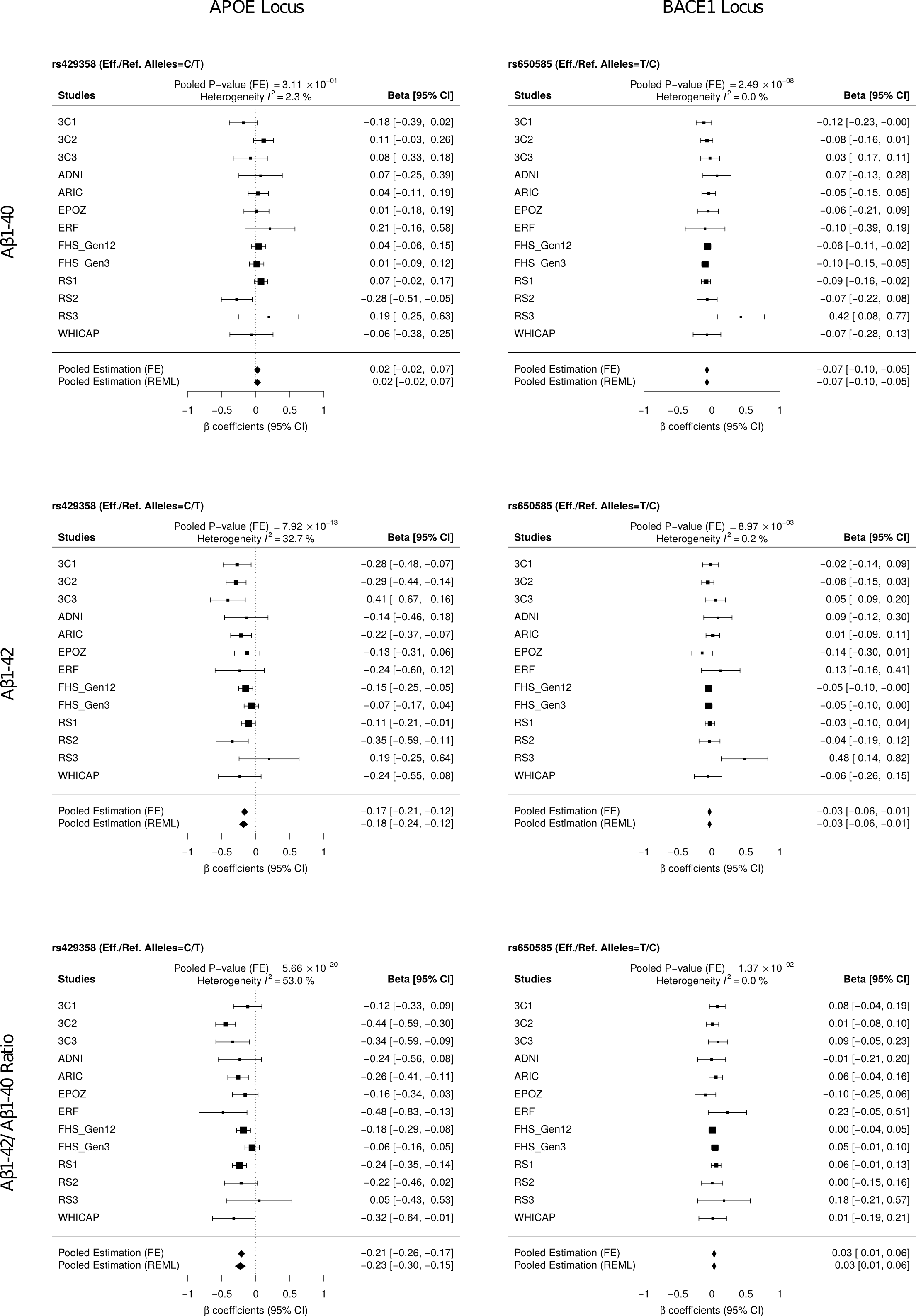

Supplementary Figure 10. Association of genotyped APOEε alleles with plasma Aβ levels

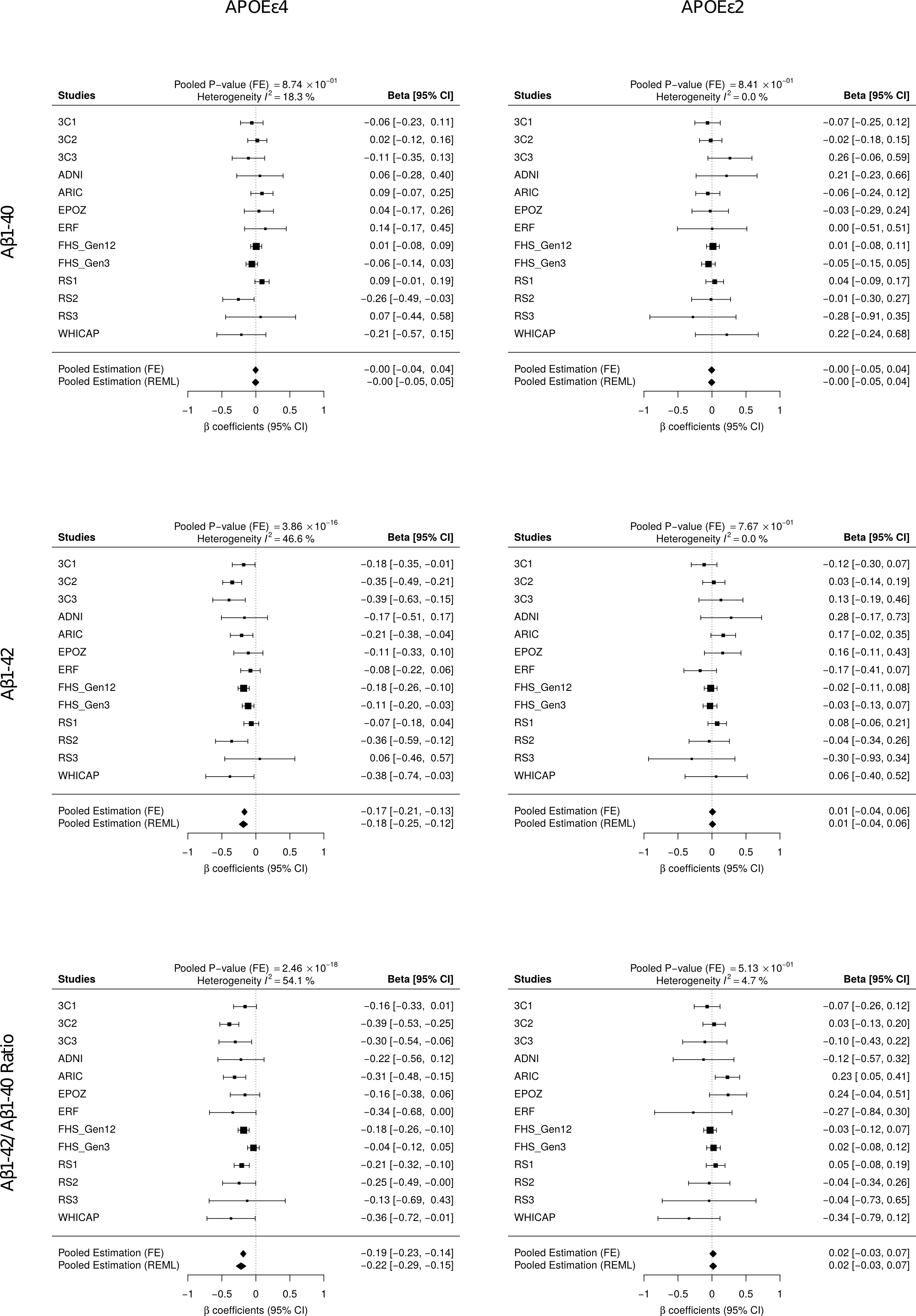

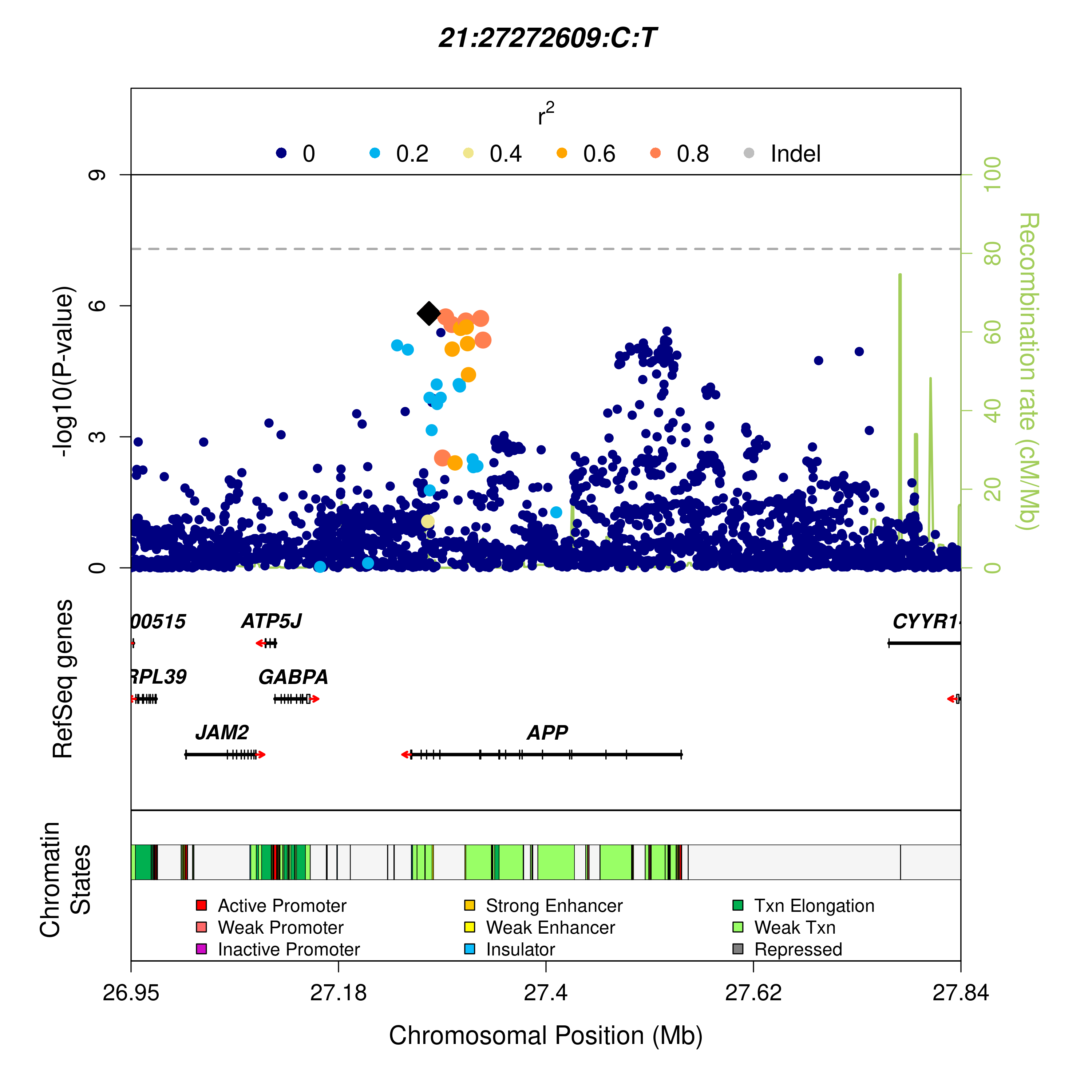
Supplementary Figure 11. Locus zoom of the APP region depicting Aβ1-40 levels association results.

Highlighted variants are the same as variants highlighted on Supplementary Figure 13.

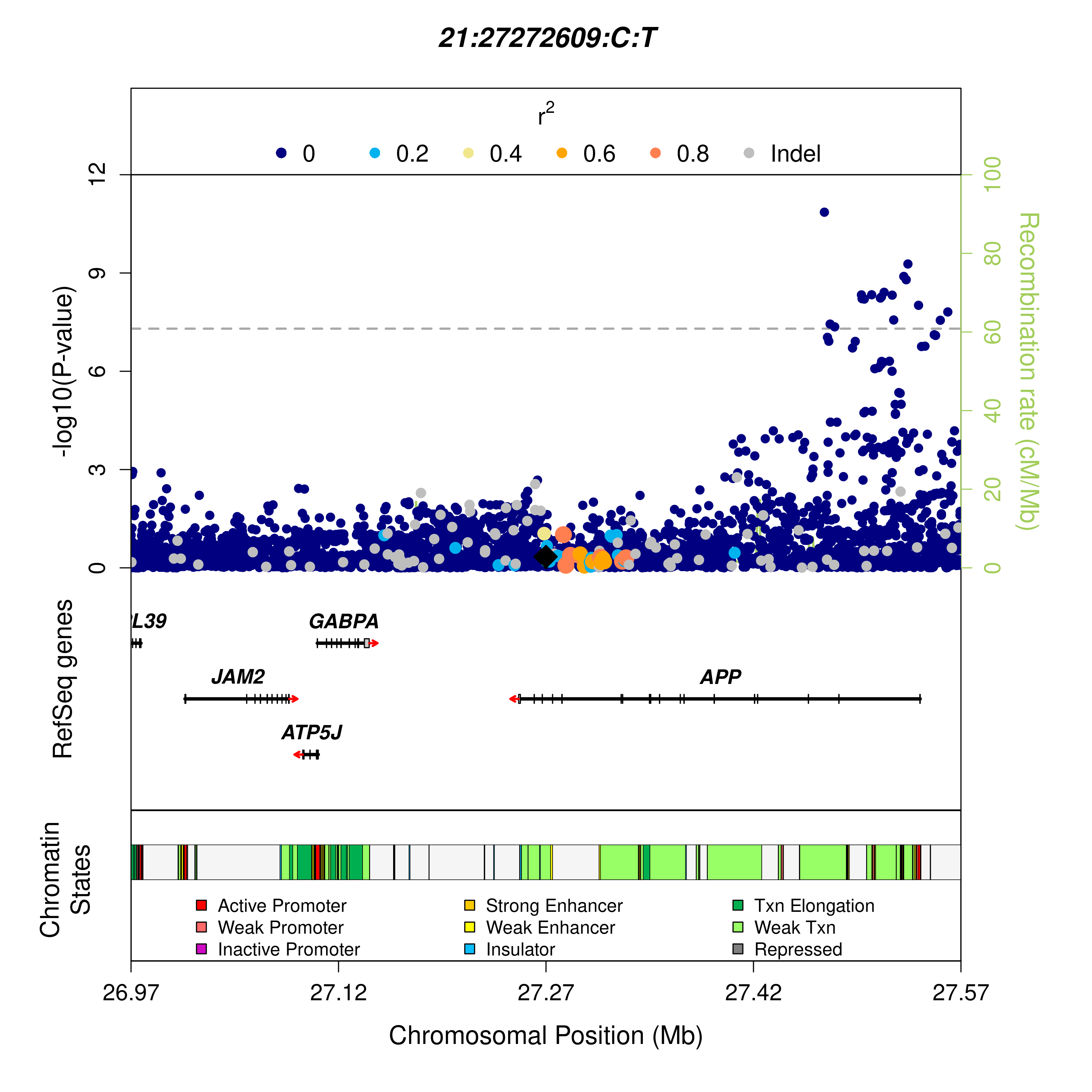
Supplementary Figure 12. Locus zoom of the APP region depicting AD association results.

Highlighted variants are the same as variants highlighted on Supplementary Figure 12.

Results were generated from [de Rojas et al., 2019].
