## Supplementary Tables Part1 for "Plasma amyloid β levels are driven by genetic variants near *APOE, BACE1, APP, PSEN2:* A genome-wide association study in over 12,000 non-demented participants"

Supplementary Table 1. Characteristics of the study populations at baseline.

| Study (N) | age, mean +/-sd | Male, % (N) | plasma Aβ1-42 pg/mL  mean +/- sd | plasma Aβ1-40 pg/mL  mean +/- sd | plasma Aβ1-42/Aβ1-40 ratio  mean +/- sd |
| --- | --- | --- | --- | --- | --- |
| 3C Batch1 (N=581) | 74.34 +/- 5.51 | 38% (221) | 38.87 +/-10.37 | 233.67 +/- 47.58 | 0.17 +/-0.05 |
| 3C Batch2 (N=1003) | 72.37 +/- 4.20 | 38% (384) | 38.56 +/- 8.53 | 192.94 +/- 45.48 | 0.21 +/-0.05 |
| 3C Batch3 (N=370) | 72.27 +/- 3.95 | 41% (150) | 34.97 +/- 7.73 | 164.16 +/- 42.61 | 0.22 +/-0.06 |
| ADNI (N=173) | 75.75 +/- 4.88 | 57% (99) | 38.05 +/-10.66 | 150.95 +/- 48.50 | 0.27 +/-0.08 |
| ARIC (N=830) | 77.18 +/- 5.30) | 40% (330) | 39.08 +/- 10.82 | 240.22 +/- 68.26 | 0.17 +/- 0.06 |
| ERF (N=129) | 64.42 +/- 4.57 | 52% (67) | 41.66 +/- 12.97 | 178.61 +/- 38.3 | 0.24 +/- 0.07 |
| EPOZ (N=397) | 70.63 +/- 6.43 | 44% (177) | 19.84 +/- 7.32 | 194.20 +/- 51.47 | 0.10 +/- 0.026 |
| FHS Gen1&2 (N=3523) | 64.33 +/-11.12 | 44% (1564) | 43.83 +/-10.23 | 160.35 +/- 37.16 | 0.28 +/-0.09 |
| FHS Gen3 (N=3212) | 47.02 +/- 8.75 | 47% (1515) | 42.74 +/- 9.94 | 243.38 +/- 54.65 | 0.18 +/-0.06 |
| RS1 (N=1549) | 68.54 +/- 8.61 | 39% (611) | 18.6 +/- 5.67 | 198.12 +/- 52.63 | 0.095 +/- 0.022 |
| RS2 (N=339) | 74.13 +/- 7.56 | 52% (176) | 18.64 +/- 9.89 | 209.58 +/- 65.14 | 0.088 +/- 0.031 |
| RS3 (N=70) | 77.23 +/- 8.28 | 50% (35) | 20.44 +/- 7.19 | 227.57 +/- 67.85 | 0.091 +/- 0.024 |
| WHICAP (N=193) | 79.89 +/- 5.54 | 40% (77) | 33.53 +/- 11.30 | 144.80 +/- 43.87 | 0.064 +/- 0.064 |

Supplementary Table 2.

See excel spreadsheet

Supplementary Table 3.

See excel spreadsheet

Supplementary Table 4.

See excel spreadsheet

Supplementary Table 5. Association results of SNP rs650585 in the BACE1 region with PET Aβ deposition.

|  | **FLR** | **rs650585** |
| --- | --- | --- |
| ***Gen 3 individuals (N=210)*** | | |
|  | Log FLR | NS |
|  | High vs Low FLR | NS |
| ***Gen 3 & APOEε4 negative individuals (N=155)*** | | |
|  | Log FLR | NS |
|  | High vs Low FLR | NS |
| ***Gen 3 & APOEε4 positive individuals (N=48)*** | | |
|  | Log FLR | β > 0 / p=0.020 |
|  | High vs Low FLR | OR=6.31 / p=0.065 |

All tests are adjusted for age, age squared and sex.

FLR : Mesure of ^11^C PiB retention in the frontal, lateral temporal and retrosplenial cortices. A threshold of 1.36 was used to define high versus low FLR. Effects were only reported when p<0.1.

NS : Non significant, OR : Odds Ratio
